# Metaplastic reinforcement of long-term potentiation in rat hippocampal area CA2 by cholinergic receptor activation

**DOI:** 10.1101/2020.10.27.358275

**Authors:** Amrita Benoy, Mohammad Zaki Bin Ibrahim, Thomas Behnisch, Sreedharan Sajikumar

**Author notes:** **Corresponding Author:** Dr Sreedharan Sajikumar, Department of Physiology, Yong Loo Lin School of Medicine, National University of Singapore, Singapore 117 597.

## Abstract

Hippocampal CA2, an inconspicuously positioned area between the well-studied CA1 and CA3 subfields, has captured research interest in recent years due to its role in the formation of social memory. The effects of synaptic depression for subsequent long-term potentiation (LTP) of synaptic transmission at entorhinal cortical (EC)-CA2 and Schaffer collateral (SC)-CA2 synapses have not been previously explored. Here we show that cholinergic receptor activation with the non-selective cholinergic agonist carbachol (CCh) triggers a long-term synaptic depression (CCh-LTD) of field excitatory postsynaptic potentials at EC- and SC-CA2 synapses in the hippocampus of adult rats. The activation of muscarinic acetylcholine receptors (mAChRs) is critical for the induction of an early phase (<100 min) of CCh-LTD, with a strong dependency upon M3 mAChR activation and a weaker one by M1 mAChRs. Interestingly, muscarinic M2 and nicotinic receptor activation are crucially involved in the late phase (>100 min) of CCh-LTD. Importantly, CCh priming lowers the threshold, in a protein synthesis-dependent manner, for the late maintenance of LTP that can be subsequently induced by high-frequency electrical stimulation at EC- or SC-CA2 pathways. The results demonstrate that CA2 synaptic learning rules are modified in a metaplastic manner, wherein synaptic modifications triggered by cholinergic stimulation can dictate the outcome of future plasticity events. Moreover, the observed enabling of late LTP at EC inputs to CA2 following the priming stimulus co-exists with concurrent sustained CCh-LTD at SC-CA2 and is dynamically scaled by modulation of SC-CA2 synaptic transmission.

**Significance Statement:** The release of the neuromodulator acetylcholine is critically involved in processes of hippocampus-dependent memory formation. Cholinergic afferents originating in the medial septum and diagonal bands of Broca terminating in the hippocampal area CA2 might play an important role in the modulation of area-specific synaptic plasticity. Our findings demonstrate that cholinergic receptor activation induces a long-term depression of synaptic transmission at entorhinal cortical- and Schaffer collateral-CA2 synapses. This cholinergic activation-mediated long-term depression displays a bidirectional metaplastic switch to long-term potentiation on a future timescale. This suggests that such bidirectional synaptic modifications triggered by the dynamic modulation of tonic cholinergic receptor activation may support the formation of CA2-dependent memories given the increased hippocampal cholinergic tone during active wakefulness observed in exploratory behaviour.

## Introduction

Neuromodulation exerts a profound influence on the modification of neural responses and synaptic plasticity (1), and allows for the adaptation and reconfiguration of neuronal circuits in response to changing external stimuli and transitions in behavioural states (2). Neuromodulatory systems in the brain influence how attention and experience regulate hippocampal representational stability and hippocampal memory formation (3-5). Activity-dependent modifications in synaptic strength such as long-term potentiation (LTP) or long-term depression (LTD) create highly coupled neuronal ensembles in the hippocampus that encode information and form memories, and are regulated by the action of neuromodulators (1). The hippocampal subfield CA2, positioned between CA3 and CA1 areas, is being increasingly considered a modulatory gateway in the hippocampal circuitry (6) with enriched expression of receptors for neuromodulators such as oxytocin (7) and vasopressin (8, 9), and receiving afferent projections from several distinct neuromodulator releasing nuclei, such as the substance P releasing supramammillary nucleus (SuM) (10-12), serotonergic median raphe nucleus, vasopressinergic paraventricular nucleus (13) and cholinergic medial septum (MS) and diagonal bands of Broca (DB) (13-16). CA2 also forms reciprocal connections projecting back to the SuM (13) and to the cholinergic nuclei of the MS-DB complex (17). The CA2 area, despite its small dimensions, is well integrated within the cortico-hippocampal network. There is direct innervation from entorhinal cortex layer II (EC LII) neurons (EC LII→CA2) at the distal CA2 dendritic region that typically expresses robust activity-dependent LTP, bypassing the indirect EC LII→dentate gyrus (DG)→CA3→CA2 pathway (18-20). In addition, CA2 neurons have strong excitatory synaptic contacts with CA1 pyramidal neurons suggestive of a functional EC LII→CA2→CA1 disynaptic loop (18). In contrast, the intrahippocampal connections from CA3 Schaffer collateral (SC) inputs that synapse onto the CA2 proximal dendritic region do not express canonical activity-dependent LTP induced by conventional high-frequency stimulation that is effective at CA3→CA1 synapses (18, 21). This distinctive resistance to plasticity displayed by CA3→CA2 synapses has been attributed to multiple suppressive factors that include the enriched expression of the scaffolding protein, regulator of G protein signaling (RGS)14 in CA2 neurons (22, 23), higher endogenous calcium buffering and enhanced calcium extrusion at CA2 dendritic spines (24), dense expression of the specialized extracellular matrix structures called perineuronal nets (25) and the relatively high expression of group III metabotropic glutamate receptors in area CA2 (26). What is striking about the action of neuromodulators on CA2 neurons is that they can potentially release this brake on plasticity at CA2 synapses (6). Substance P (27) and agonists to the vasopressin 1b (Avpr1b) and oxytocin (Oxtr) receptors (28), for instance, can trigger synaptic potentiation at the plasticity-resistant CA3 Schaffer collateral (SC)-CA2 synapses.

Among the hippocampal CA subfields, CA2 *stratum pyramidale* has been shown to have the highest density of the acetylcholine-synthesizing enzyme, choline acetyltransferase (ChAT)-positive punctate immunoprecipitates in the human hippocampal formation (29). ChAT-immunoreactivity is also found to be more concentrated in the CA2 and CA3 regions of the rat hippocampus (30, 31). The CA2 region also shows the highest expression of acetylcholinesterase (AChE) enzyme relative to other CA regions (32, 33). In view of the reciprocal connectivity of area CA2 with the cholinergic medial septum and diagonal bands of Broca and the enriched expression of cholinergic marker enzymes, ChAT and AChE, in CA2 relative to other CA regions, we hypothesized that cholinergic modulation shapes synaptic plasticity in hippocampal CA2 neurons.

In the present study, we have investigated how synaptic plasticity is regulated at hippocampal CA2 synapses through the activation of cholinergic receptors with the non-selective cholinergic agonist carbachol (CCh). We observed a long-lasting synaptic depression at both entorhinal cortical and Schaffer collateral inputs to CA2 neurons upon activation of cholinergic receptors. This effect ‘primes’ CA2 synapses for future long-term potentiation in a metaplastic manner. Metaplasticity refers to a higher-order form of synaptic plasticity in which activity-dependent modifications in synaptic strength at a given point in time are not only an outcome of the timing and frequency of presynaptic stimulation and postsynaptic activity, but also are dependent on the previous history of synaptic activity (34, 35). The data presented in this study demonstrate that the metaplastic trigger of cholinergic receptor activation-induced LTD is protein synthesis-dependent and enables late LTP at EC-CA2 synapses subjected to weak tetanizing stimuli subthreshold for the expression of lasting LTP. More intriguingly, the plasticity-resistant SC-CA2 synapses are also primed for late LTP in a similar fashion, with the difference being that LTP is maintained only when CCh-LTD is followed by a strong tetanization that is otherwise ineffective for LTP expression at SC-CA2. Interestingly, the enabling of late LTP at EC-CA2 synapses co-exists with concomitant CCh-induced synaptic depression at SC-CA2, and is dynamically down-scaled by the modulation of SC-CA2 synaptic transmission, alluding to a heterosynaptic interplay between afferent Schaffer collateral stimulation and the metaplastic reinforcement of LTP at cortical inputs to CA2 neurons. Together, this study brings to light a previously unrecognized role for metaplasticity by cholinergic activation in the dynamic temporal regulation of synaptic efficacy in CA2 neurons, which could potentially mediate experience-dependent and behavioural state-dependent modulation of synaptic plasticity, and thereby learning and memory.

## Results

### Cholinergic receptor activation triggers protein synthesis-dependent and NMDAR-independent long-term depression at both entorhinal cortical- and Schaffer collateral-CA2 synapses

On the basis of our hypothesis that cholinergic modulation could shape synaptic plasticity in hippocampal CA2 neurons, we first sought to investigate changes in synaptic efficacy along the proximo-distal dendritic axis of neurons in area CA2 owing to cholinergic receptor activation. To this end, we performed extracellular field potential recordings in acute rat hippocampal slices, that recorded synaptic activity from a population of CA2 neurons as field excitatory post synaptic potentials (fEPSPs) from the distal entorhinal cortical (EC)-CA2 and proximal Schaffer collateral (SC)-CA2 synapses. Control recordings were performed at basal stimulation frequency (0.2 Hz) that resulted in stable synaptic responses for 5 h after a baseline of 30 min at EC-CA2 and SC-CA2 synapses (Fig. 1*B*, EC-CA2, red circles, Wilcox test, *P* > 0.05; SC-CA2, blue circles, Wilcox test, *P* > 0.05 at any given time point), thus demonstrating the stability of the field potential recordings performed in this study. To probe the effect of cholinergic stimulation in CA2 neurons, the non-selective cholinergic receptor agonist, carbachol (CCh) (50 µM) was bath-applied for 30 min after taking a baseline recording of 30 min (Fig. 1*C*) resulting in a sustained synaptic depression (CCh-LTD) that lasted for 4 h post CCh application at both EC-CA2 (Fig. 1*C*, red circles, 270 min: Wilcox test, *P* = 0.0020) and SC-CA2 synapses (Fig. 1*C*, blue circles, 270 min: Wilcox test, *P* = 0.0039). A previous study has shown that activity-dependent LTD in response to low-frequency electrical stimulation exhibits a heterogeneous expression pattern in CA2 neurons in the rat hippocampus (21). The cumulative frequency distribution of fEPSP (%) values from Fig. 1*C*, 60 min and 270 min into CCh application, is shown in Fig. 1*D* and 1*E*, respectively. The data indicate some form of heterogeneity, at a minor level, for CCh-LTD displayed by EC- and SC-CA2 synapses, wherein CA2 synapses in a few experiments displayed a reversal to baseline values over time, particularly evident at SC-CA2 synapses 270 min into CCh application (Fig. 1*E*). However, in the majority of experiments, the EC- and SC-CA2 synapses remained significantly depressed upon cholinergic activation by CCh application. We also noted that the cumulative frequency distribution of field potentials between EC and SC inputs to CA2 at 60 min and 270 min into CCh application are not significantly different (Fig. 1*D*, Kolmogorov-Smirnov test, *P* > 0.05; Fig. 1*E*, Kolmogorov-Smirnov test, *P* > 0.05). Next, we sought to explore how the CCh-induced depression at synapses in area CA2 compares to that in area CA1. To determine this, we recorded synaptic responses from the hippocampal CA1 region upon independently stimulating CA1 *stratum radiatum* (SR) where Schaffer collaterals terminate on CA1 neurons (SC-CA1), and CA1 *stratum lacunosum-moleculare* (SLM) where axons primarily from entorhinal cortical layer III terminate on CA1 neurons (EC-CA1) (Fig. S1*A*). Briefly, CCh (50 µM) was bath-applied for a duration of 30 min after a 30 min stable baseline and synaptic responses from SC-CA1 and EC-CA1 were continued to be recorded up to 4 h post CCh application at basal stimulation frequency throughout as shown in Fig. S1*B*. We observed that, similar to SC-CA2 synapses, synaptic responses in CA1 SR (SC-CA1) displayed sustained CCh-induced synaptic depression up to 4 h post CCh application (Fig. S1*B*, blue circles, 270 min: Wilcox test, *P* = 0.0059). However, unlike EC-CA2 that remained significantly depressed up to 270 min into CCh application, synaptic responses at CA1 SLM (EC-CA1) reversed to its baseline levels by 210 min into CCh application (Fig. S1*B*, yellow circles, 210 min: Wilcox test, *P* > 0.05). The CCh-induced depression has also been previously reported to be expressed differently between the SR and SLM of CA1 area in that CA1 SR displays greater CCh-induced synaptic depression compared to CA1 SLM (36). The effect of CCh on synaptic responses in SR and SLM of CA2 area (data from Fig. 1*C*) and CA1 area (data from Fig. S1*B*) is compared against each other at 30 min, 60 min, 180 min and 270 min into CCh application in Fig. S1*D*. Together, the summarized data in Fig. S1*D* suggest that while the maintenance of LTD in response to cholinergic receptor activation differs between CA2 SLM (EC-CA2) and CA1 SLM (EC-CA1) (Fig. S1*D*, 180 min: *U* test, *P* = 0.0232, 270 min: *U* test, *P* = 0.0115), it is similar between CA2 SR (SC-CA2) and CA1 SR (SC-CA1) (Fig. S1*D*, 180 min: *U* test, *P* > 0.05, 270 min: *U* test, *P* > 0.05), and the CCh-induced immediate depression is significantly greater at SC-CA1 compared to SC-CA2 synapses (Fig. S1*D*, 30 min: *U* test, *P* = 0.0052), while it does not significantly vary between EC-CA1 and EC-CA2 synapses (Fig. S1*D*, 30 min: *U* test, *P* > 0.05).

**Fig. 1.**
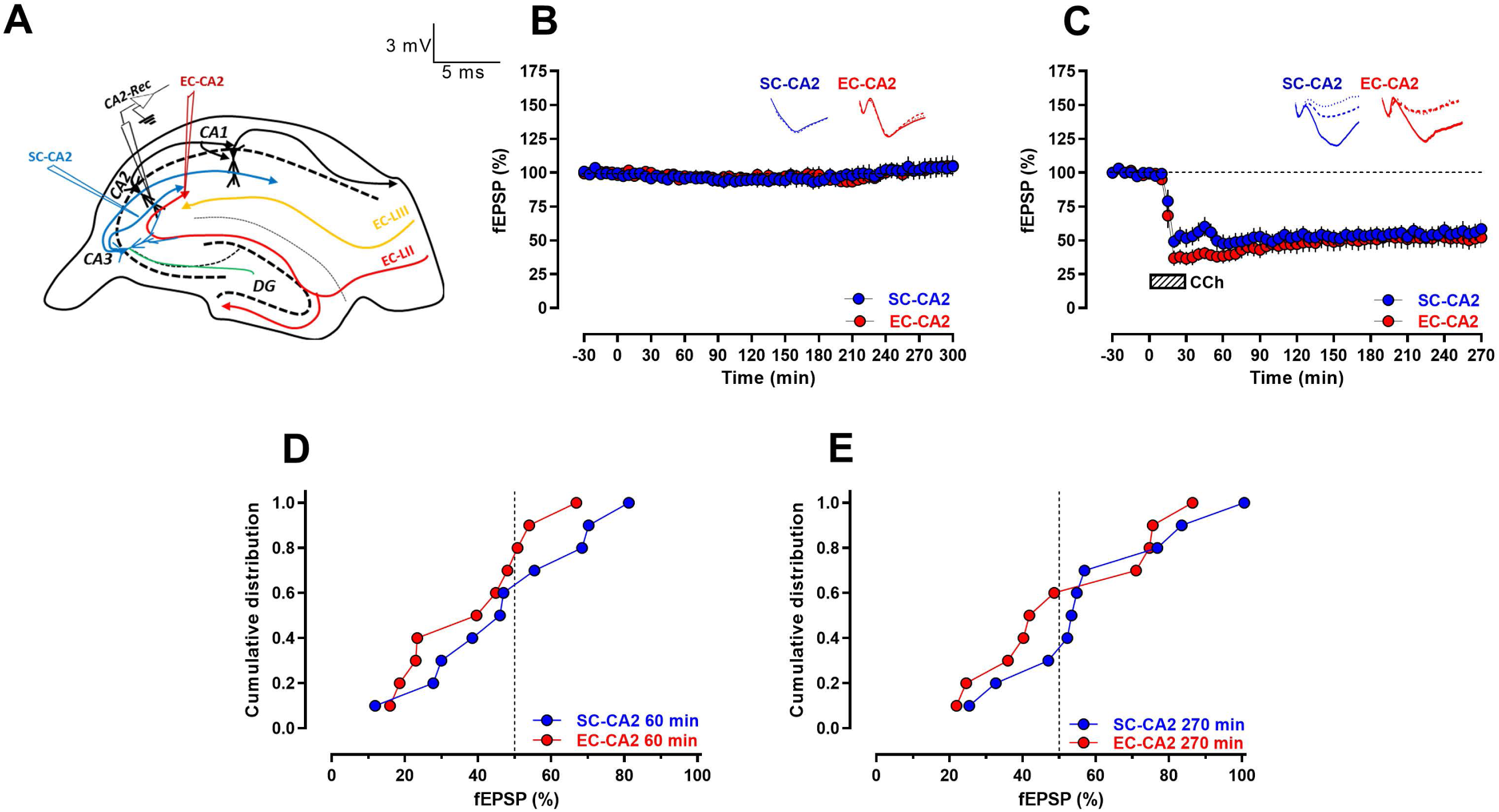
Non-selective cholinergic receptor agonist carbachol (CCh) triggers long-lasting synaptic depression at entorhinal cortical and Schaffer collateral synaptic inputs to CA2 neurons. *(A)* A schematic diagram depicting the location of electrodes in the acute hippocampal slices for the extracellular field electrophysiology experiments performed in this study. Blue inverted triangle represents the placement of electrode in the *stratum radiatum* stimulating the Schaffer collateral (SC) afferents to CA2. Red inverted triangle represents the location of electrode in the *stratum lacunosum-moleculare* stimulating the entorhinal cortical (EC) afferents to CA2. Black inverted triangle depicts the position of the recording electrode in the distal CA2 dendritic region. *(B)* Control experiments at basal synaptic stimulation frequency indicate the long-lasting stability up to 5 h of SC-CA2 (blue circles) and EC-CA2 (red circles) fEPSP recordings in this study (*n* = 6). *(C)* Bath application of 50 µM CCh for 30 min after a 30 min stable baseline induced a long-lasting synaptic depression (CCh-LTD) at SC and EC inputs to CA2 at basal synaptic stimulation, lasting for 4 h post CCh application (*n* = 10). All data show mean ± SEM. Analog traces depict representative SC-CA2 (blue) and EC-CA2 (red) fEPSPs within the initial 30 min baseline recording (closed line) before CCh application, and 60 min (dotted line) and 270 min (hatched line) after baseline. *(D-E)* Summary data with cumulative frequency distribution plots of SC-CA2 and EC-CA2 fEPSP(%) values from Fig. 1*C* at 60 min *(D)* and 270 min *(E)* into CCh application. (Scale bars for analog traces in *B-C*: 3 mV/5 ms.)

Further, we analysed the paired pulse ratio at both EC-CA2 (Fig. S2*A-B*) and SC-CA2 (Fig. S2*C-D*) before CCh application during the 30 min baseline recording (‘control’) and 60 min into CCh application (‘CCh’) to determine whether CCh-LTD has a presynaptic or a postsynaptic locus of expression. Pairs of stimulation pulses with an interstimulus interval of 10 ms, 30 ms, 50 ms, 70 ms, 100 ms, 150 ms and 250 ms were delivered before CCh application and 60 min into CCh application. We observed that for 10 ms interstimulus interval, the paired pulse ratio at EC-CA2 60 min into CCh application is significantly higher than control paired pulse ratio before CCh application (Fig. S2*A*, Wilcox test, *P* = 0.0156). The percent paired pulse ratio at 10 ms interstimulus interval in EC-CA2 normalized by the average control paired pulse ratio is represented in Fig. S2*B*, that shows a significant increase of 34.1 ± 0.317 % for 60 min into CCh application. The observed increase in paired pulse ratio suggests that LTD at EC-CA2 60 min into CCh application involves a reduction in the probability presynaptic glutamate release. However, the paired pulse ratio at SC-CA2 was not significantly different before CCh and 60 min into CCh application (Fig. S2*C*, Wilcox test, *P* > 0.05 at 10 ms, 30 ms, 50 ms, 70 ms, 100 ms, 150 ms and 250 ms interstimulus intervals). The percent paired pulse ratio at 10 ms interstimulus interval in SC-CA2 normalized by the average control paired pulse ratio is represented in Fig. S2*D* that shows no significant change between control and CCh groups. This suggests that, unlike EC-CA2, the synaptic depression at SC-CA2 60 min into CCh application is not associated with a significant change in the presynaptic glutamate release probability. The synaptic depression from CCh application was also shown to be protein synthesis-dependent as bath-application of the protein synthesis inhibitor emetine (20 µM) significantly attenuated the maintenance of synaptic depression at CA2 synapses in response to CCh (Fig. S3*A*). As shown in Fig. S3*A*, emetine was applied for a total of 1 h (30 min prior to co-application with CCh for another 30 min) and this resulted in fEPSPs at EC- and SC-CA2 synapses gradually returning to levels not significantly different from their respective baseline recordings by 265 min into emetine application for EC-CA2 (Fig. S3*A*, red circles, 265 min: Wilcox test, *P >* 0.05) and by 240 min into emetine application for SC-CA2 (Fig. S3*A*, blue circles, 240 min: Wilcox test, *P >* 0.05). Fig. S3*B* represents the fEPSP(%) values from Fig. S3*A* (CCh + emetine) and Fig. 1*C* (CCh alone), at 60 min and 270 min into CCh application, compared against each other. Although the acute CCh-LTD (60 min) is comparable with and without emetine (Fig. S3*A* and Fig. 1*C*; *U* test, *P* > 0.05 for EC-CA2 and SC-CA2), the CCh-LTD has decayed back to baseline by 4.5 h (270 min) in the presence of emetine at both EC (*U* test, *P* = 0.0097) and SC (*U* test, *P* = 0.0185) inputs to CA2 (Fig. S3*B*).

The LTD induced by CCh, however, was not NMDA receptor-dependent as co-application of the NMDAR antagonist D-AP5 (50 µM) with CCh did not attenuate CCh-LTD at either synaptic inputs to CA2 with fEPSPs significantly lower than baseline levels even by 270 min into CCh application (Fig. S3*C*, EC-CA2, red circles, 270 min: Wilcox test, *P* = 0.0078; SC-CA2, blue circles, 270 min: Wilcox test, *P* = 0.0234). Fig. S3*D* presents a comparative analysis of fEPSP(%) from Fig. S3*C* (CCh + D-AP5) and Fig. 1*C* (CCh alone), at 60 min and 270 min into CCh application, and shows that fEPSPs for both acute CCh-LTD (60 min) and sustained CCh-LTD (270 min) in Fig. 1*C* and S3*C* are not significantly different from each other (Fig. S3*D, U* test, *P >* 0.05). We further determined that CCh-LTD at CA2 synapses was independent of continuous afferent synaptic drive as we observed a significant and lasting LTD at EC- and SC-CA2 synapses even when test stimulation was suspended for 1 h (30 min during application of CCh and 30 min immediately following CCh application) (Fig. S3*E*). As demonstrated in Fig. S3*E*, both EC- and SC-CA2 displayed significant synaptic depression compared to their respective baseline recordings, despite the 1 h pause in test stimulation (EC-CA2, red circles, 270 min: Wilcox test, *P* = 0.0313; SC-CA2, blue circles, 270 min: Wilcox test, *P* = 0.0313). Fig. S3*F* compares fEPSP(%) from Fig. S3*E* (CCh + suspension of test stimulus) and Fig. 1*C* (CCh alone), at 60 min and 270 min into CCh application, and shows that fEPSPs for both acute CCh-LTD (60 min) and sustained CCh-LTD (270 min) in Fig. 1*C* and S3*E* are not significantly different from each other (Fig. S3*F, U* test, *P >* 0.05).

### Differential role for muscarinic and nicotinic acetylcholine receptors in facilitating CCh-LTD at CA2 synapses

Acetylcholine (ACh) exerts its effects on neural activity by acting on ionotropic nicotinic ACh receptors (nAChRs) and metabotropic muscarinic ACh receptors (mAChRs) (37). Given the expression of cholinergic receptors belonging to both ACh receptor classes in the CA2 region (38-40), we were intrigued to understand the relative roles played by the different cholinergic receptors in mediating CCh-LTD at CA2 synapses. Interestingly, application of the selective muscarinic M1 receptor antagonist, pirenzepine (0.5 µM), for a total duration of 50 min (20 min prior to co-application with CCh for another 30 min) resulted in fEPSPs at EC-CA2 and SC-CA2 returning to levels not significantly different from their individual baseline values by 90 min into pirenzepine application (Fig. 2*A*, EC-CA2, red circles, 90 min: Wilcox test, *P* > 0.05; SC-CA2, blue circles, 90 min: Wilcox test, *P* > 0.05). Application of the selective muscarinic M2 receptor antagonist, AF-DX 116 (2 µM) for a total duration of 50 min (20 min prior to co-application with CCh for 30 min) abrogated the late maintenance of CCh-LTD at EC-CA2 synapses with fEPSPs gradually returning to levels not significantly different from its baseline values by 200 min into the M2 antagonist application (Fig. 2*B*, EC-CA2, red circles, 200 min: Wilcox test, *P* > 0.05). In contrast, at SC-CA2, the M2 receptor antagonist did not abolish the late maintenance of CCh-LTD and showed a significant CCh-induced synaptic depression even by 270 min into application of the antagonist (Fig. 2*B*, SC-CA2, blue circles, 270 min: Wilcox test, *P* = 0.0098). However, when compared to SC-CA2 synaptic depression from CCh alone in Fig. 1*C*, there was a significant attenuation of CCh-LTD at SC-CA2 with the M2 antagonist in Fig. 2*B* (SC-CA2, 250 min into CCh application: *U* test, *P* = 0.0241), suggesting a role for muscarinic M2 receptor activation in CCh-LTD at SC-CA2, nonetheless. However, the observed difference in CCh-LTD attenuation between EC-CA2 and SC-CA2 by M2 receptor blockade suggests that M2 receptor activation plays a greater role in mediating CCh-LTD at EC-CA2 compared to SC-CA2 synapses. Interestingly, application of the muscarinic M3 receptor antagonist, 4-DAMP (1 µM), for a total duration of 50 min (20 min prior to co-application with CCh for 30 min) as shown in Fig. 2*C*, significantly attenuated the induction itself of CCh-LTD when compared to application of CCh alone depicted in Fig. 1*C* (EC-CA2, 30 min into CCh application: *U* test, *P* < 0.0001; SC-CA2, 30 min into CCh application: *U* test, *P* = 0.0021). Muscarinic M3 receptor blockade with 4-DAMP, when compared to M1 and M2 receptor blockade, also resulted in faster decay of CCh-induced CA2 synaptic depression, with fEPSPs reverting to levels not significantly different from its baseline values by 70 min into 4-DAMP application at EC-CA2 and SC-CA2 synapses (Fig. 2*C*, EC-CA2, red circles, 70 min: Wilcox test, *P* > 0.05; SC-CA2, blue circles, 70 min: Wilcox test, *P* > 0.05). Nevertheless, the complete blockade of muscarinic receptors with the non-selective muscarinic receptor antagonist, atropine (1 µM) bath-applied for a total duration of 50 min (20 min prior to and 30 min co-application with CCh), resulted in the induction of CCh-LTD being fully abolished, with fEPSPs at EC-CA2 and SC-CA2 not significantly different from their respective baseline levels at any given time point into the application of atropine (Fig. 2*D*, EC-CA2, red circles, Wilcox test, *P* > 0.05; SC-CA2, blue circles, Wilcox test, *P* > 0.05). This highlights a critical role for muscarinic receptor activation in the induction of CCh-LTD at CA2 synapses. In contrast, application of the non-selective nicotinic receptor antagonist, mecamylamine (20 µM) for 50 min (20 min prior to and 30 min co-application with CCh) resulted in the blockade of CCh-LTD only by 265 min into mecamylamine application at EC-CA2 (Fig. 2*E*, EC-CA2, red circles, 265 min: Wilcox test, *P* > 0.05) and by 225 min into mecamylamine application at SC-CA2 (Fig. 2*E*, SC-CA2, blue circles, 225 min: Wilcox test, *P* > 0.05). The time points at which CCh-LTD is abrogated in Fig. 2*A, B, C* and *E*, suggest that M3 receptor blockade attenuates CCh-LTD faster than by M1, M2 or nicotinic receptor blockade, implying a prominent role for muscarinic M3 receptor activation in the early facilitation of CCh-LTD followed by M1 receptors, and suggest that activation of muscarinic M2 and nicotinic receptors plays a role in the late maintenance of CCh-LTD.

**Fig. 2.**
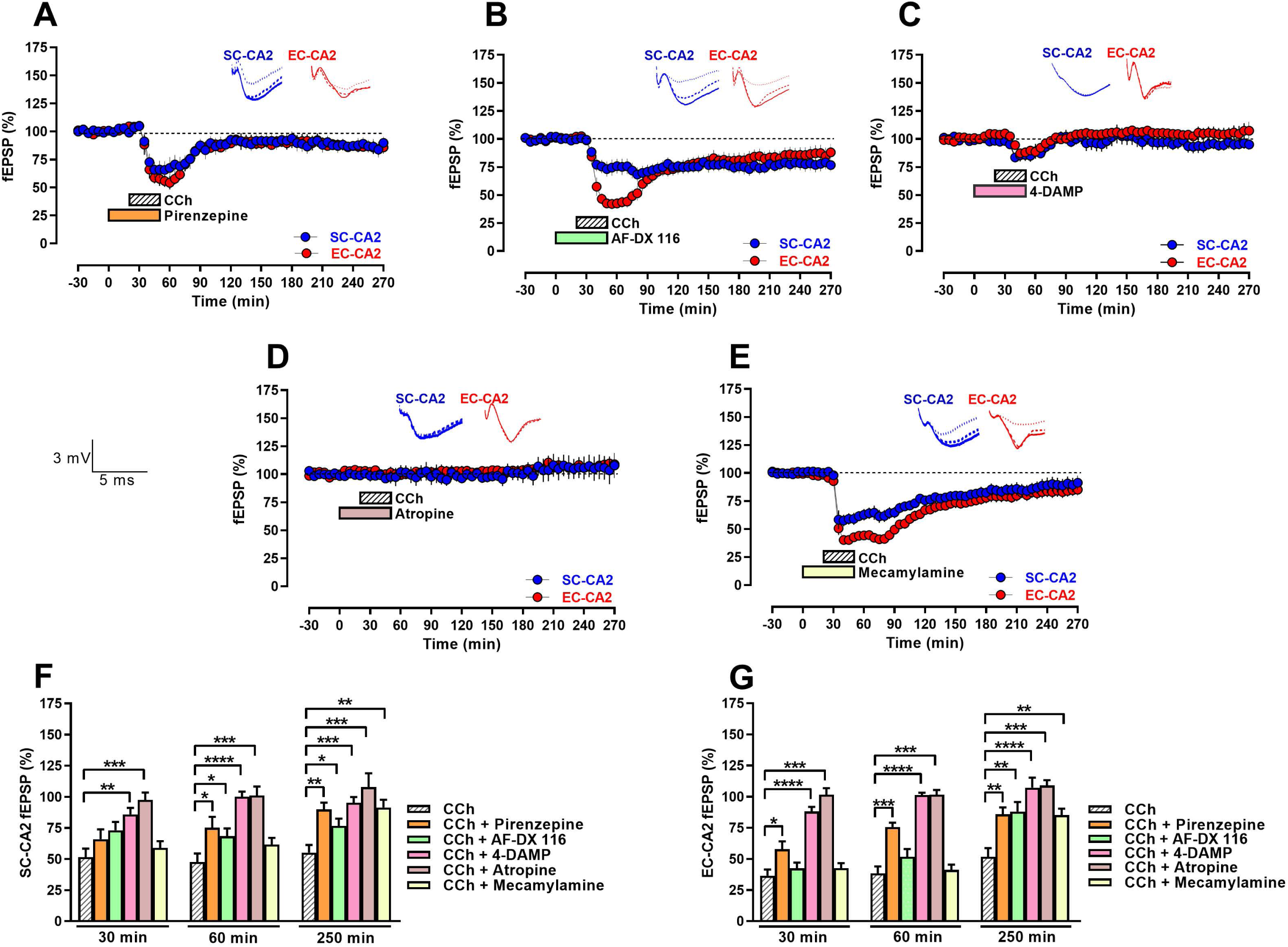
Muscarinic and nicotinic acetylcholine receptor activation play varied roles in the induction, early and late phase of CCh-LTD at EC-CA2 and SC-CA2 synapses. *(A)* Muscarinic M1 receptor antagonist, pirenzepine (0.5 µM) is bath-applied for a total of 50 min (20 min before CCh and 30 min co-application with CCh) following a baseline of 30 min, resulting in the blockade of LTD by 70 min into CCh application at both EC-CA2 and SC-CA2 synapses (*n* = 7). *(B)* Muscarinic M2 receptor antagonist, AF-DX 116 (2 µM) is bath-applied for a total of 50 min (20 min before CCh and 30 min co-application with CCh) following a baseline of 30 min, resulting in the blockade of LTD by 180 min into CCh application at EC-CA2, while significant depression persists even into the late phase at SC-CA2 (*n* = 11). *(C)* Muscarinic M3 receptor antagonist, 4-DAMP (1 µM) is bath-applied for a total of 50 min (20 min before CCh and 30 min co-application with CCh) following a baseline of 30 min, resulting in only a brief synaptic depression that is abrogated by 50 min into CCh application at both EC- and SC-CA2 (*n* = 8), faster than M1 and M2 receptors. *(D)* The non-selective muscarinic receptor antagonist, atropine (1 µM) when bath-applied for a total of 50 min (20 min before CCh and 30 min co-application with CCh) following a baseline of 30 min, results in the complete abrogation of CCh-LTD at both EC- and SC-CA2 (*n* = 6). *(E)* The non-selective nicotinic receptor antagonist, mecamylamine (20 µM) when bath-applied for a total of 50 min (20 min before CCh and 30 min co-application with CCh) following a baseline of 30 min, however, abrogates only the very late maintenance of CCh-LTD by 245 min into CCh application at EC-CA2 and by 205 min into CCh application at SC-CA2 (*n* = 12). *(F-G)* Summary bar graphs comparing fEPSP(%) between Fig. 1*C* (CCh alone) and Fig. 2*A-E* (CCh + various cholinergic receptor antagonists) at 30 min, 60 min and 250 min into CCh application for SC-CA2 *(F)* and EC-CA2 *(G)* synapses, outline the varied roles for the different cholinergic receptors in the induction, early and late phases of CCh-LTD at SC- and EC-CA2. All data show mean ± SEM. Significant differences between groups are indicated as * *P* < 0.05, ** *P* < 0.01, *\*\*\* P* < 0.001, *\*\*\*\* P* < 0.0001. Analog traces depict representative EC-CA2 (red) and SC-CA2 (blue) fEPSPs within 30 min before CCh application (closed line), at 60 min into CCh application (dotted line) and at 250 min into CCh application (hatched line). (Scale bars for analog traces in *A-E*: 3 mV/5 ms.)

The observed role for muscarinic M2 and nicotinic receptors is in contrast to the recent study in mice where M2 and nicotinic receptors were not observed to be involved in CCh-induced synaptic depression in CA2 neurons (41). However, we suspect that the aforementioned study could not detect a role for M2 and nicotinic receptors in CCh-LTD due to a short recording period. Similar to their results, we also observed that the acute depression following CCh application was not blocked by M2 or nicotinic antagonists. A comparative analysis of fEPSP(%) from Fig. 2*A-E* (CCh + various cholinergic receptor antagonists) with Fig. 1*C* (CCh alone), at 30 min, 60 min and 250 min into CCh application, at SC-CA2 and EC-CA2 synapses is presented in Fig. 2*F* and Fig. 2*G*, respectively. Together, the data presented in Fig. 2*A-G* indicate that while activation of nicotinic receptors and muscarinic M1, M2 and M3 receptors facilitate CCh-LTD, albeit with differences in their contribution to the induction, early and late phases of CCh-LTD, the induction itself of CCh-LTD at CA2 synapses is mediated through muscarinic and not nicotinic receptor activation.

### Prior synaptic depression mediated by cholinergic receptor activation primes CA2 synapses for long-term potentiation

As the maintenance of CCh-induced synaptic depression at CA2 synapses was shown to be protein synthesis-dependent, we explored if such protein synthesis and/or other activated signaling downstream of cholinergic receptor activation could alter the state of CA2 synapses to regulate subsequently induced synaptic plasticity. We hypothesized that if there is a common pool of plasticity-related proteins between LTD and LTP, then this could potentially prime the depressed synapses for LTP as well when stimulus strength is altered. To this end, we performed single-pathway experiments wherein we stimulated the EC inputs alone to CA2 at the *stratum lacunosum-moleculare* and recorded fEPSPs from the CA2 dendritic region. Upon recording a 30 min stable baseline, CCh was bath-applied for 30 min, which resulted in a significant synaptic depression at the EC-CA2 synapses (Fig. 3*A*, red circles, 60 min: Wilcox test, *P* = 0.0078). However, after 60 min into CCh application, the stimulus strength of the EC input was increased such that the fEPSPs returned to values comparable to the initial average baseline values before CCh application (Fig. 3*A*, red circles, 80 min: *U* test, *P* > 0.05). As shown in Fig. 3*A*, a new stable baseline was thereafter recorded at the increased stimulus strength for another 30 min, following which a weak tetanization (WTET) was delivered to the EC-CA2 pathway. We observed that priming from CCh-LTD results in the transformation of a transient early LTP (E-LTP) to a long-lasting late LTP (L-LTP) at EC-CA2 maintained for 3 h post WTET (Fig. 3*A*, red circles, 275 min: Wilcox test, *P* = 0.0078). In the absence of CCh priming, weak tetanic stimulation at EC-CA2 normally results only in a transient synaptic potentiation that is short lasting (E-LTP) and decays to baseline levels by 95 min post WTET (Fig. 3*B*, red circles, 95 min: Wilcox test, *P* > 0.05). Thus, CCh-LTD serves as a metaplastic trigger for the reinforcement of subsequently induced LTP by stimuli subthreshold for producing L-LTP, at EC-CA2 synapses. In order to understand if this ‘primed’ LTP is protein synthesis-dependent, we bath-applied the protein synthesis inhibitor, emetine 15 min into resetting the baseline by increasing the EC input stimulus strength after 60 min into CCh application (Fig. 3*C*). As shown in Fig. 3*C*, after recording a new stable baseline for 15 min at the increased stimulus strength, emetine was bath-applied for a total duration of 60 min, and WTET was delivered to EC-CA2 30 min into emetine application. However, a significant synaptic potentiation was still observed in EC-CA2 3 h post WTET (Fig. 3*C*, red circles, 290 min: Wilcox test, *P* = 0.0039). This prompted us to explore if early protein synthesis inhibition during CCh-LTD could affect the primed LTP. Interestingly, when emetine was applied for a total duration of 60 min (30 min prior to and 30 min co-application with CCh) as shown in Fig. 3*D*, the EC-CA2 synaptic potentials returned to levels not significantly different from the reset baseline potentials by 95 min post WTET (Fig. 3*D*, red circles, 220 min: Wilcox test, *P* > 0.05). Although there is a small yet significant post tetanic potentiation at EC-CA2 following WTET in Fig. 3*D* (red circles, 1 min post WTET: Wilcox test, *P* = 0.0156), it is significantly attenuated in comparison to the post tetanic potentiation in Fig. 3*A* (1 min post WTET: *U* test, *P* = 0.0006). These data demonstrate that protein synthesis initiated in response to CCh-induced synaptic depression is critical for the early induction phase as well as the enabling of LTP upon a sequential increase in stimulus strength followed by weak tetanic stimulation at EC-CA2. The bar graph in Fig. 3*E* represents EC-CA2 fEPSP(%) at 3 h post WTET from Fig. 3*A*, Fig. 3*C* and Fig. 3*D*, summarizing the effect of protein synthesis inhibition at different time points on the persistence of CCh-primed LTP at EC-CA2 synapses.

**Fig. 3.**
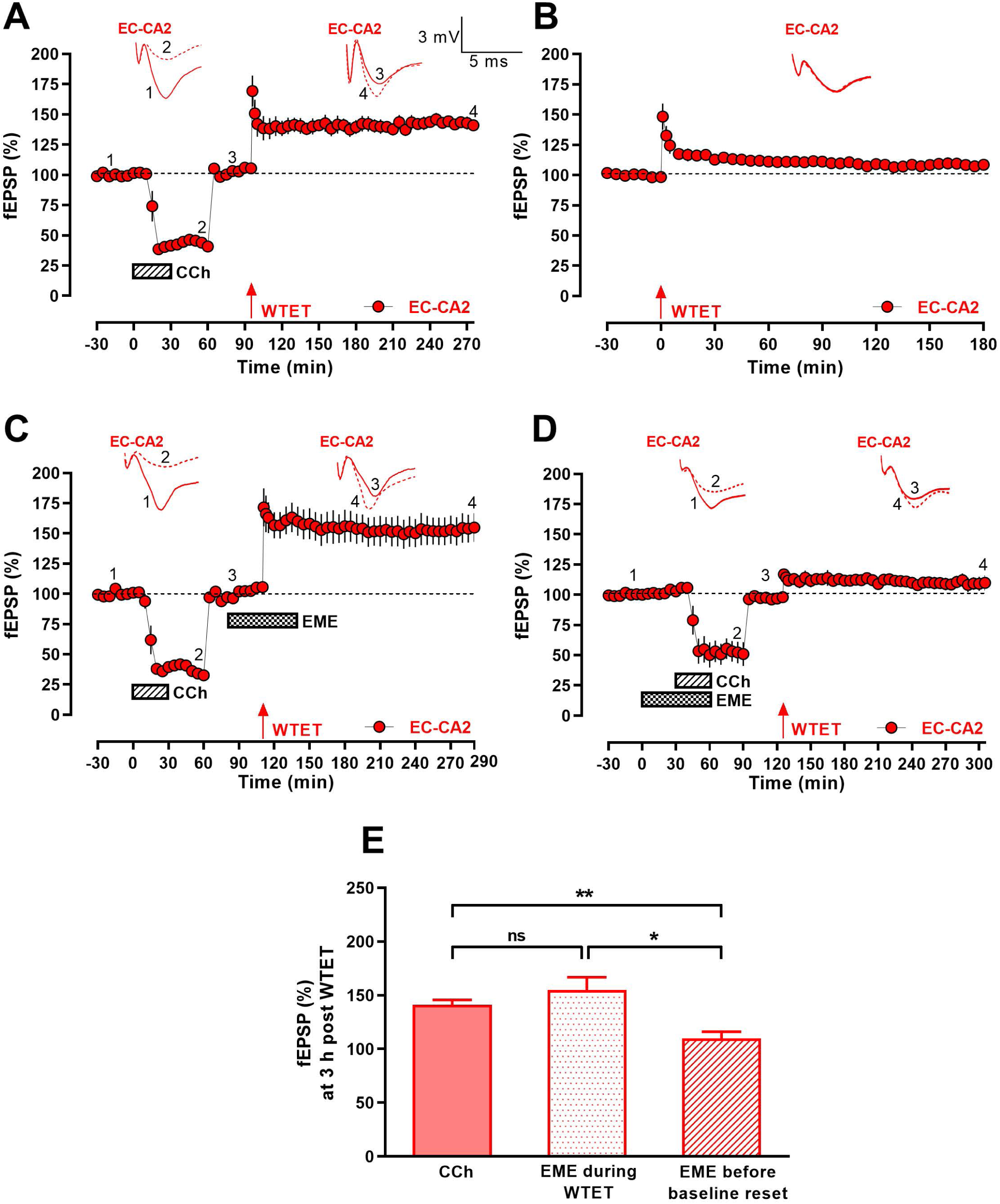
Prior synaptic depression by cholinergic receptor activation primes EC-CA2 synapses for late LTP induced by weak tetanic stimulation at a later point in time. *(A)* In single-pathway experiments where EC fibres alone to CA2 were stimulated, CCh was bath-applied for 30 min following an initial stable baseline of 30 min, resulting in a significant synaptic depression at EC-CA2. 1 h into application of CCh, the stimulus strength at EC-CA2 was increased and fEPSPs reverted to levels comparable to the initial baseline values exhibited prior to application of CCh. Following a new reset stable baseline for 30 min, weak tetanic stimulation (WTET) was delivered at EC-CA2, resulting in persistent LTP (*n* = 8). *(B)* Control single-pathway experiment without prior CCh priming, where WTET was delivered at EC-CA2 following a stable baseline of 30 min, resulted in a short-lasting potentiation termed early LTP that decayed to baseline by 95 min post WTET (*n* = 8). *(C)* Late inhibition of protein synthesis during the delivery of WTET at EC-CA2 does not abrogate the reinforcement of CCh-primed LTP. CCh was bath-applied for 30 min following an initial stable baseline of 30 min, resulting in a significant synaptic depression at EC-CA2. 1 h into application of CCh, the stimulus strength at EC-CA2 was increased and fEPSPs reverted to levels comparable to the initial baseline values exhibited prior to application of CCh. Following a new reset stable baseline of 15 min, the protein synthesis inhibitor emetine (EME; 20 µM) was applied for 1 h while WTET was delivered at EC-CA2 30 min into emetine application. However, LTP still persisted for 3 h post WTET (*n* = 9). *(D)* Early inhibition of protein synthesis during CCh-LTD prevents the reinforcement of WTET-induced LTP at EC-CA2. Emetine was bath-applied for a total duration of 60 min (30 min before CCh and 30 min co-application with CCh) after a 30 min baseline. 1 h into application of CCh, the stimulus strength at EC-CA2 was increased and fEPSPs reverted to levels comparable to the initial baseline values. 30 min into the new reset stable baseline, WTET was delivered at EC-CA2, which however, did not result in persistent potentiation at EC-CA2 (*n* = 7). *(E)* Summary bar graph comparing fEPSP (%) at 3 h post WTET at EC-CA2 from Fig. 3*A*, 3*C* and 3*D*. All data show mean ± SEM. Differences between groups are indicated as * *P* < 0.05, ** *P* < 0.01 and ns *P* > 0.05. Analog traces of Fig. 3*A*, 3*C* and 3*D* depict representative EC-CA2 (red) fEPSPs within 30 min before CCh application (closed line-1), at 60 min into CCh application (hatched line-2), within 30 min before WTET (closed line-3), and at 180 min post WTET (hatched line-4). Analog traces of Fig. 3*B* depict representative EC-CA2 fEPSPs within the 30 min baseline before WTET (closed line) and at 180 min post WTET (hatched line). (Scale bars for analog traces in *A-D*: 3 mV/5 ms.)

Similar to the LTP reinforcement observed at EC-CA2 synapses that were weakly stimulated subsequent to CCh-LTD, we next explored if this metaplastic effect is also displayed by the SC-CA2 synapses. We designed a single-pathway experiment with a timeline similar to that in Fig. 3*A* with the exception that in place of EC stimulation, the SC-CA2 pathway was stimulated at the proximal *stratum radiatum* and fEPSPs were recorded from the CA2 dendritic region (Fig. 4*A*). After 30 min of an initial stable baseline, CCh was bath-applied for 30 min which resulted in a significant synaptic depression at SC-CA2 (Fig. 4*A*, SC-CA2, blue circles, 60 min: Wilcox test, *P* = 0.0078). 60 min into CCh application, the input stimulus strength for SC-CA2 was increased such that its fEPSPs returned to values comparable to the initial average baseline values before CCh application (Fig. 4*A*, SC-CA2, blue circles, 80 min: *U* test, *P* > 0.05). After recording a new reset stable baseline for 30 min at the increased stimulus strength, WTET was delivered to the SC-CA2 pathway. However, unlike the EC-CA2 pathway, the SC-CA2 synapses did not display a persistent LTP and fEPSPs decayed to the reset baseline potentials by 15 min post WTET (Fig. 4*A*, SC-CA2, blue circles, 110 min: Wilcox test, *P >* 0.05). Fig. 4*B* shows the control single-pathway experiment without CCh priming where a WTET delivered at SC-CA2 following a 30 min stable baseline recording results in a transient potentiation that decays to baseline levels by 15 min post WTET (Fig. 4*B*, SC-CA2, blue circles, 15 min: Wilcox test, *P* > 0.05). As SC-CA2 synapses are different than EC-CA2 in their plasticity expression and have been shown in multiple previous studies to be resistant to canonical activity-dependent LTP (18, 21, 27), we were intrigued to explore if a stronger stimulus could facilitate LTP reinforcement after CCh priming in the SC-CA2 pathway. To test this possibility, a strong tetanization (STET) consisting of three trains of high-frequency stimulation at 100 Hz was delivered at SC-CA2 following CCh priming (Fig. 4*C*). Similar to the experiment timeline in Fig. 4*A*, after 30 min of an initial stable baseline, CCh was bath-applied for 30 min which resulted in a significant synaptic depression at SC-CA2 (Fig. 4*C*, SC-CA2, blue circles, 60 min: Wilcox test, *P* = 0.0005). 60 min into CCh application, the input stimulus strength for SC-CA2 was increased to revert fEPSPs to values comparable to the initial average baseline values before CCh application (Fig. 4*C*, SC-CA2, blue circles, 80 min: *U* test, *P >* 0.05). Following a new stable baseline of 30 min, STET was delivered to the SC-CA2 pathway, which resulted in L-LTP that remained significantly higher than its reset baseline potentials for 3 h (Fig. 4*C*, SC-CA2, blue circles, 275 min: Wilcox test, *P* = 0.0398). Fig. 4*D* shows the single-pathway control recording where SC-CA2 synapses were subjected to STET following a 30 min stable baseline recording, resulting in only a transient potentiation that quickly decayed to baseline levels by 15 min post STET (Fig. 4*D*, SC-CA2, blue circles, 15 min: Wilcox test, *P* > 0.05). Thus, cholinergic receptor activation-induced synaptic depression primes both EC and SC synapses onto CA2 for long-term potentiation, albeit in response to different strength of plasticity-inducing tetanic stimuli.

**Fig. 4.**
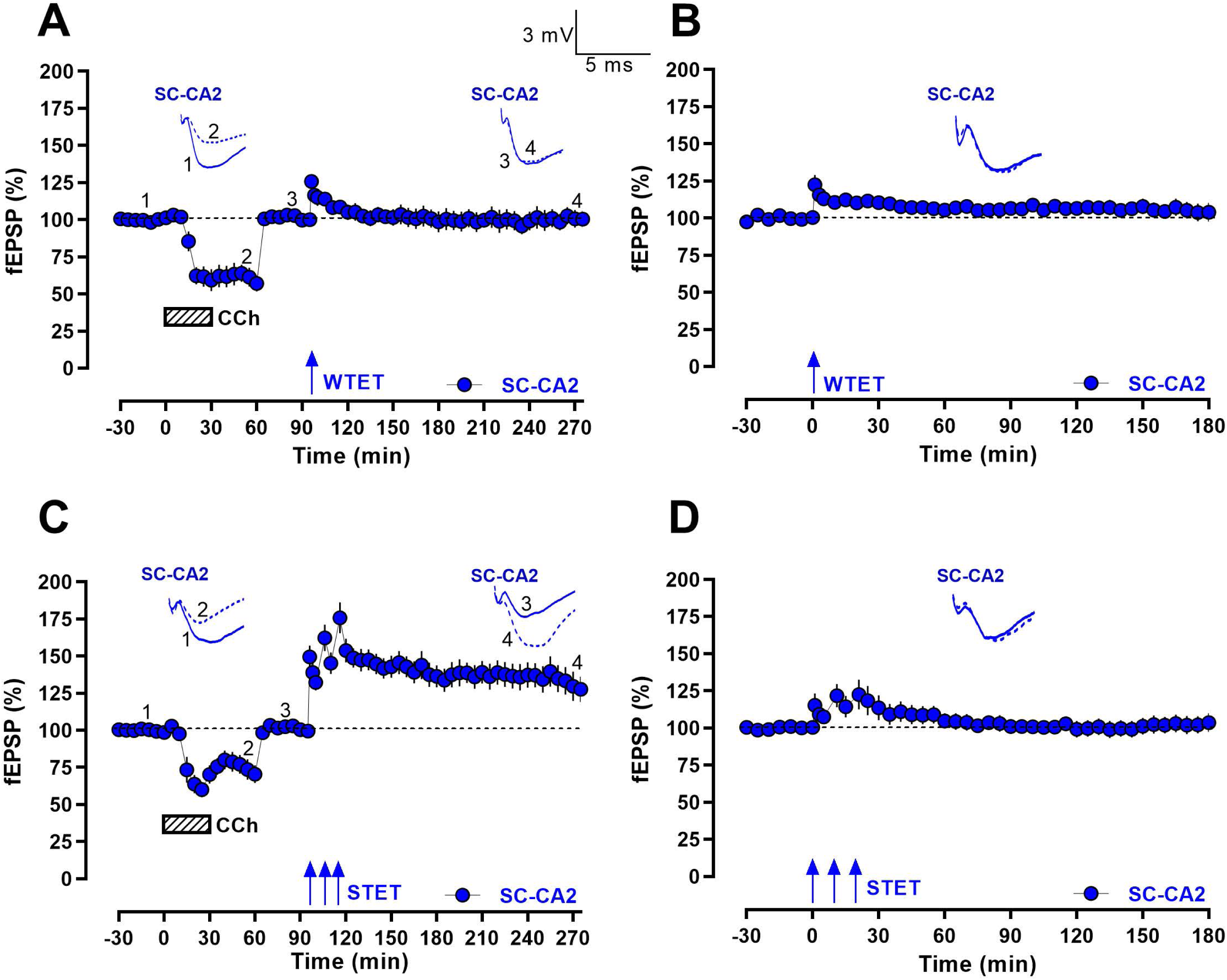
Late LTP is expressed at SC-CA2 synapses in response to strong but not weak tetanic stimulation subsequent to CCh-LTD. *(A)* In single-pathway experiments where SC fibres to CA2 alone were stimulated, CCh was bath-applied for 30 min following an initial stable baseline of 30 min, resulting in a significant synaptic depression at SC-CA2. 1 h into application of CCh, the stimulus strength at SC-CA2 was increased and fEPSPs reverted to levels comparable to the initial baseline values exhibited prior to CCh application. Following a new reset stable baseline for 30 min, weak tetanizing stimuli (WTET) was delivered at SC-CA2, which elicited only a transient potentiation that quickly decayed to baseline levels by 15 min post WTET (*n* = 8). *(B)* Control single-pathway experiment without prior CCh priming, where WTET is delivered at SC-CA2 following a stable baseline of 30 min, resulting in a similar level of transient potentiation as Fig. 4*A* that decays to baseline levels by 15 min post WTET (*n* = 6). *(C)* Strong tetanic stimulation (STET) following CCh-LTD elicits LTP reinforcement at SC-CA2. CCh was bath-applied for 30 min following an initial stable baseline of 30 min, resulting in a significant synaptic depression at SC-CA2. 1 h into CCh application, the stimulus strength at SC-CA2 was increased and fEPSPs reverted to levels comparable to the initial baseline values exhibited prior to application of CCh. Following a new reset stable baseline for 30 min, STET was delivered at SC-CA2, which results in synaptic potentiation reinforced for up to 3 h (*n* = 13). *(D)* Control single-pathway experiment without prior CCh priming, where STET was delivered at SC-CA2 following a stable baseline of 30 min, resulted in only a transient potentiation that quickly decayed to baseline levels by 15 min post STET, thus demonstrating the resistance displayed by SC-CA2 synapses to activity-dependent LTP in the absence of CCh priming (*n* = 10). All data show mean ± SEM. Analog traces of Fig. 4*A* and 4*C* depict representative SC-CA2 (blue) fEPSPs within 30 min before CCh application (closed line-1), at 60 min into CCh application (hatched line-2), within 30 min before TET (closed line-3), and at 180 min post TET (hatched line-4). Analog traces of Fig. 4*B* and 4*D* depict representative SC-CA2 fEPSPs within the 30 min baseline before TET (closed line) and at 180 min post TET (hatched line). (Scale bars for analog traces in *A-D*: 3 mV/5 ms.)

### Reinforcement of CCh-primed LTP at cortical synaptic inputs to CA2 co-exists with concurrent and sustained CCh-LTD in the Schaffer collateral pathway

The spatial segregation of incoming information via the SC and EC pathways converging onto CA2 proximal and distal dendritic regions, respectively, raises the possibility of a heterosynaptic trans-compartmental interaction in the processing of incoming synaptic inputs to CA2 neurons. With the existence of a primed LTP at EC-CA2 synapses demonstrated in response to prior cholinergic activation-induced synaptic depression in single-pathway experiments (Fig. 3*A*), we were thus intrigued to perform two-pathway experiments with similar experimental design to explore if there is an interaction between the EC and SC pathways modulating the expression of the observed LTP. To determine this, we performed experiments wherein we stimulated both EC and SC fibres onto CA2, but differentially changed the stimulus strength of both pathways after CCh-induced synaptic depression. First, we performed a two-pathway experiment involving stimulation of EC and SC fibres, wherein a stable baseline was recorded for 30 min, following which CCh was bath-applied for 30 min which resulted in a significant synaptic depression at EC (Fig. 5*A*, red circles, 60 min: Wilcox test, *P* = 0.0156) and SC (Fig. 5*A*, blue circles, 60 min: Wilcox test, *P* = 0.0156) inputs to CA2. After 60 min into CCh application, the stimulus strength of EC inputs alone was increased such that fEPSPs at EC-CA2 reverted to values comparable to its initial average baseline values before CCh application (Fig. 5*A*, EC-CA2, red circles, 80 min: *U* test, *P* > 0.05). Stimulus strength for the SC-CA2 pathway was unaltered such that SC-CA2 synapses displayed significant and sustained CCh-LTD until the end of the recording period (Fig. 5*A*, SC-CA2, blue circles, 275 min: Wilcox test, *P* = 0.0156). In the EC-CA2 pathway, however, after 30 min of a stable reset baseline, WTET was delivered, which resulted in L-LTP lasting 3 h (Fig. 5*A*, EC-CA2, red circles, 275 min: Wilcox test, *P* = 0.0156). Fig. 5*B* shows the control two-pathway experiment without CCh priming, where WTET is delivered to EC-CA2 following a 30 min stable baseline recording, resulting in only a short-lasting E-LTP that decays to baseline levels by 40 min post WTET (Fig. 5*B*, EC-CA2, red circles, 40 min: Wilcox test, *P >* 0.05). SC-CA2 synapses are subjected to baseline stimulation throughout and display stable synaptic responses until the end of the recording period (Fig. 5*B*, SC-CA2, blue circles, Wilcox test, *P* > 0.05 at any given time point). As shown in Fig. 5*C*, we also compared changes in the presynaptic release probability, by performing paired pulse stimulation at EC-CA2 synapses at three different points in the experiment timeline (before CCh application, at a time point in the reset baseline post CCh application and after 30 min post WTET), for two-pathway experiments performed from Fig. 5*A*. Pairs of stimulation pulses with an interstimulus interval of 10 ms, 30 ms, 50 ms, 70 ms, 100 ms, 150 ms and 250 ms were delivered at EC-CA2 at a point during baseline recording before CCh application (‘pre CCh’), at a point during the new reset baseline taken after increasing stimulus strength at EC (‘post CCh baseline reset’) and after 30 min post WTET (‘post WTET’). However, the paired pulse ratios at EC-CA2 between the three experimental time points were not significantly different from each other (Fig. 5*C*, Kruskal-Wallis test with Dunn’s multiple comparisons *post hoc* analysis, *P* > 0.05 at 10 ms, 30 ms, 50 ms, 70 ms, 100 ms, 150 ms and 250 ms interstimulus intervals). Fig. 5*D* shows the percent paired pulse ratio normalized by the average ‘pre CCh’ paired pulse ratio for the paired pulse stimulation experiments at 10 ms interstimulus interval from Fig. 5*C*. Although there is a slight increase of 24.2 ± 0.082 % at 10 ms interstimulus interval between the normalized EC-CA2 paired pulse ratio before CCh application (‘pre CCh’) and at a point in the new reset baseline after increasing EC stimulus strength (‘post CCh baseline reset’), the difference is not statistically significant (Fig. 5*D*, Kruskal-Wallis test with Dunn’s multiple comparisons *post hoc* analysis, *P* > 0.05). Similarly, the normalized EC-CA2 paired pulse ratio at a point in the new reset baseline (‘post CCh baseline reset’) and after 30 min post WTET (‘post WTET’) are not significantly different from each other (Fig. 5*D*, Kruskal-Wallis test with Dunn’s multiple comparisons *post hoc* analysis, *P* > 0.05), suggesting that the observed CCh-primed LTP at EC-CA2 in the two-pathway experiment is not associated with a significant change in the presynaptic glutamate release probability after 30 min post WTET. However, comparing Fig. 5*D* and Fig. S2*B*, we noted that after resetting to a new stable baseline upon increasing stimulus strength, the paired pulse ratio at EC-CA2 is not significantly different from the paired pulse ratio before CCh application, while we had observed a significant increase in paired pulse ratio (10 ms interstimulus interval) 1 h into CCh application at EC-CA2 (Fig. S2*B*). This shows that the decrease in presynaptic release probability at EC-CA2 seen 1 h into CCh application before resetting the baseline, is abrogated and reverted to a level comparable to what was before CCh application when the baseline is reset.

**Fig. 5.**
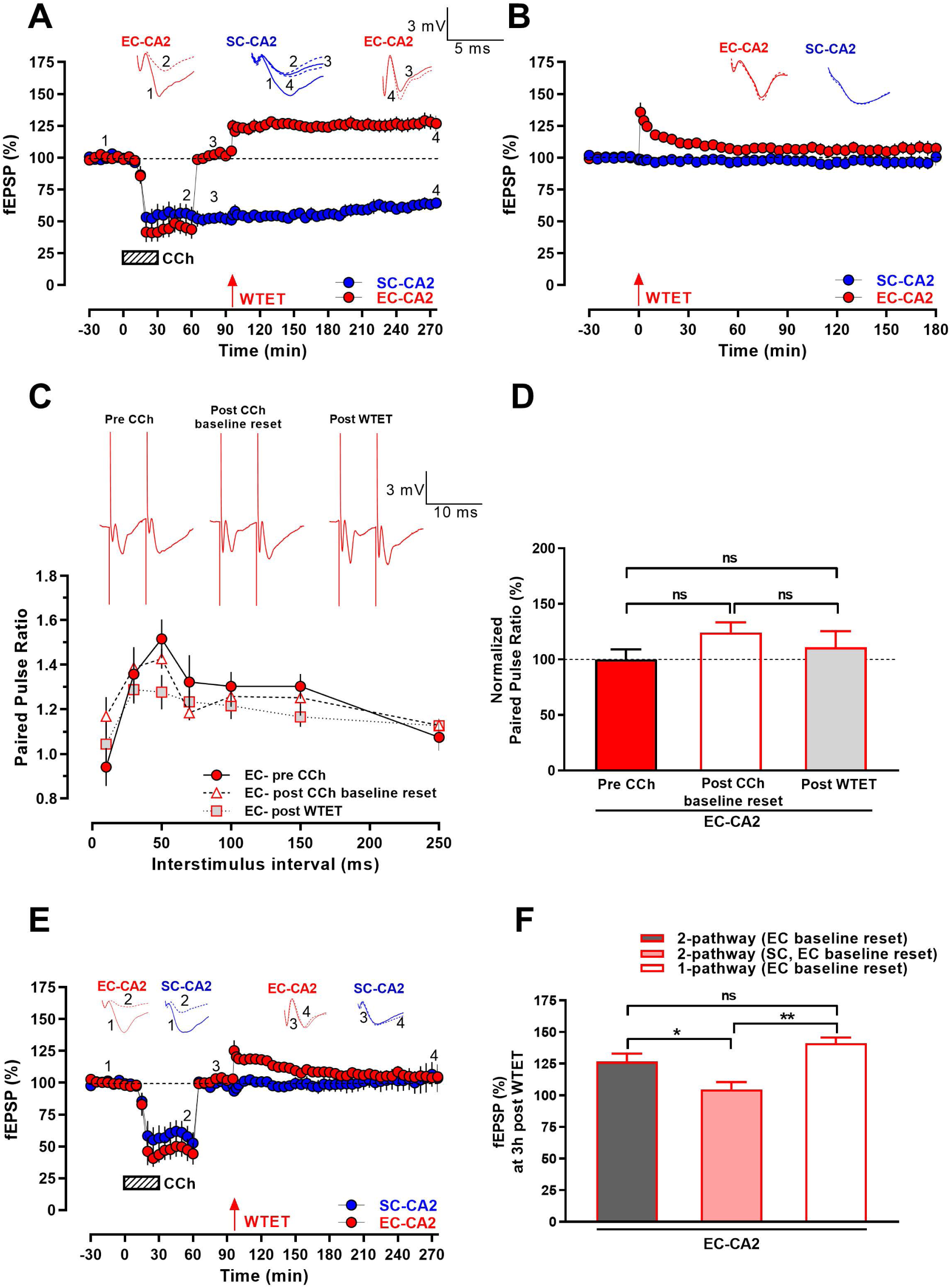
CCh-primed late LTP at EC-CA2 steadily co-exists with concurrent CCh-LTD at SC-CA2 synapses. *(A)* In two-pathway experiments where EC and SC fibres to CA2 were stimulated, CCh was bath-applied for 30 min following an initial stable baseline of 30 min, resulting in a significant synaptic depression at EC-CA2 and SC-CA2. 1 h into application of CCh, the stimulus strength at EC-CA2 alone was increased and EC-CA2 fEPSPs reverted to levels comparable to the initial baseline values exhibited prior to CCh application, while stimulus strength at SC-CA2 was unaltered. Following a new reset stable baseline at EC-CA2 for 30 min, weak tetanic stimulus (WTET) was delivered at EC-CA2, resulting in persistent expression of LTP at EC-CA2, that co-exists with sustained CCh-LTD displayed by SC-CA2 synapses (*n* = 7). *(B)* Control two-pathway experiment without prior CCh priming, where WTET was delivered at EC-CA2 following a stable baseline of 30 min, resulting in only a short-lasting early LTP that decayed to baseline by 40 min post WTET while SC-CA2 synapses displayed stable baseline synaptic responses until the end of the recording period (*n* = 7). *(C)* Paired pulse ratio at EC-CA2 was analysed for the experiments in Fig. 5*A*, at three stages during the recording: In the baseline prior to CCh application (‘pre CCh’), in the reset baseline post increasing stimulus strength at EC-CA2 (‘post CCh baseline reset’), and 30 min post WTET at EC-CA2 (‘post WTET’). Paired pulse ratios for stimulation pulses delivered to EC-CA2 at interstimulus intervals of 10 ms, 30 ms, 50 ms, 70 ms, 100 ms, 150 ms and 250 ms, do not display significant differences between ‘Pre CCh’, ‘Post CCh baseline reset’ and ‘Post WTET’ groups (*n*= 6). *(D)* EC-CA2 paired pulse ratios at 10 ms interstimulus interval from Fig. 5*C* were normalized by the average ‘Pre CCh’ paired pulse ratio and compared amongst the three stages. No significant change is detected in the normalized paired pulse ratio (%) between the ‘Pre CCh’, ‘Post CCh baseline reset’ and ‘Post WTET’ groups. *(E)* Increasing stimulus strength at SC-CA2 along with EC-CA2 synapses following CCh priming prevents LTP reinforcement at EC-CA2 subsequently subjected to WTET. In two-pathway experiments where EC and SC fibres to CA2 were stimulated, CCh was bath-applied for 30 min following an initial stable baseline of 30 min, resulting in a significant synaptic depression at EC-CA2 and SC-CA2. 1 h into CCh application, the stimulus strength at both EC-CA2 and SC-CA2 was increased such that EC-CA2 and SC-CA2 fEPSPs revert to levels comparable to the initial baseline values exhibited prior to CCh application. Following a new reset stable baseline at EC-CA2 and SC-CA2 for 30 min, WTET was delivered at EC-CA2 alone. Unlike Fig. 5*A*, only a short-lasting early LTP ensues at EC-CA2 that decays to baseline levels by 35 min post WTET, while SC-CA2 synapses display stable synaptic responses at the reset baseline levels until the end of the recording period (*n* = 6). *(F)* Summary bar graph comparing fEPSP(%) at EC-CA2 3 h post WTET between Fig. 3*A* [‘1-pathway (EC baseline reset)’], Fig. 5*A* [‘2-pathway (EC baseline reset)’] and Fig. 5*E* [‘2-pathway (SC, EC baseline reset)’], indicating comparable levels of LTP reinforcement between single and two-pathway experiments where stimulus strength at EC inputs alone was increased following CCh priming, and significant attenuation in fEPSP(%) post WTET when stimulus strength at SC inputs was also increased. All data show mean ± SEM. Differences between groups are indicated as * *P* < 0.05, ** *P* < 0.01 and ns *P* > 0.05. Analog traces of Fig. 5*A* and 5*E* depict representative EC-CA2 (red) and SC-CA2 (blue) fEPSPs within 30 min before CCh application (closed line-1), at 60 min into CCh application (hatched line-2), within 30 min before WTET at EC-CA2 (closed line-3), and at 180 min post WTET at EC-CA2 (hatched line-4). Analog traces of Fig. 5*B* depict representative EC-CA2 and SC-CA2 fEPSPs within the 30 min baseline before WTET at EC-CA2 (closed line) and at 180 min post WTET at EC-CA2 (hatched line). Analog traces of Fig. 5*C* depict representative EC-CA2 (red) fEPSPs for the paired pulse stimulation separated by a 10 ms interstimulus interval. (Scale bars for analog traces in *A, B*, and *E*: 3 mV/5 ms, in *C*: 3 mV/10 ms.)

In a separate two-pathway experiment, we followed the same experimental design as Fig. 5*A* with the exception that the stimulus strength was increased post 60 min into CCh application at SC-CA2 in addition to the EC-CA2 pathway (Fig. 5*E*). After a 30 min stable baseline, CCh was bath-applied for 30 min, which resulted in a significant synaptic depression at both EC (Fig. 5*E*, red circles, 60 min: Wilcox test, *P* = 0.0313) and SC inputs to CA2 (Fig. 5*E*, blue circles, 60 min: Wilcox test, *P* = 0.0313). After 60 min into CCh application, stimulus strength at both SC and EC pathways was increased such that synaptic potentials at both pathways reverted to values comparable to the initial average baseline values before CCh application (Fig. 5*E*, EC-CA2, red circles, 80 min: *U* test, *P* > 0.05; SC-CA2, blue circles, 80 min: *U* test, *P* > 0.05). 30 min following the reset stable baseline at EC-CA2 and SC-CA2, WTET was delivered at the EC-CA2 pathway alone (Fig. 5*E*). Unlike Fig. 5*A*, this did not result in the late persistence of LTP at EC-CA2 with fEPSPs at EC-CA2 decaying to baseline levels by 35 min post WTET (Fig. 5*E*, EC-CA2, red circles, 130 min: Wilcox test, *P* > 0.05). fEPSPs at SC-CA2 were not significantly different from its reset baseline levels until the end of the recording period (Fig. 5*E*, SC-CA2, blue circles, 275 min: Wilcox test, *P* > 0.05). Thus, the persistence of LTP at EC-CA2 following CCh-induced synaptic depression is influenced by the strength of concurrent synaptic responses at SC-CA2 and co-exists with sustained CCh-LTD in the SC-CA2 pathway. EC-CA2 postsynaptic potentials 3 h following WTET were not significantly different between the two-pathway experiment in Fig. 5*A* and the single-pathway EC-CA2 experiment in Fig. 3*A* (EC-CA2, 3 h post WTET: *U* test, *P* > 0.05). The bar graph in Fig. 5*F* summarizes the net EC-CA2 fEPSP(%) at 3 h post WTET from Fig. 5*A*, Fig. 5*E* and Fig. 3*A*. However, the net post tetanic potentiation immediately following WTET was significantly higher in the single-pathway EC-CA2 experiments in Fig. 3*A* when compared to the two-pathway experiments performed in Fig. 5*A* (EC-CA2, 1 min post WTET: *U* test, *P* = 0.0059) and Fig. 5*E* (EC-CA2, 1 min post WTET: *U* test, *P* = 0.0127). It is unlikely that the observed reduction in the magnitude of net post tetanic potentiation is due to the afferent stimulation of both pathways, as we observed comparable levels of net post tetanic potentiation at EC-CA2 in the control single-pathway (Fig. 3*B*) and two-pathway (Fig. 5*B*) EC-CA2 WTET experiments without CCh priming (EC-CA2, 1 min post WTET: *U* test, *P >* 0.05). We speculate that synaptic stimulation of SC-CA2 upon CCh priming interferes with the expression of post tetanic potentiation at EC-CA2 in the two-pathway experiment contributing to the observed scaling down of post tetanic potentiation, although the mechanisms mediating this heterosynaptic effect remain to be determined.

Further, in order to understand how prior synaptic depression induced by cholinergic receptor activation may alter the level of depolarization in CA2 neurons in response to subsequent delivery of WTET at EC-CA2 in comparison to control experiments where WTET was delivered at EC-CA2 in the absence of cholinergic activation, we analyzed and compared the tetanic stimuli mediated summation of field potentials at EC-CA2 with and without CCh priming (Fig. 6*A-C*). Fig. 6*A* shows the average EC-CA2 field potential traces in response to 100 Hz/s from single (Fig. 3*B*) and two-pathway (Fig. 5*B*) EC-CA2 WTET control experiments without CCh priming, and from single (Fig. 3*A*) and two-pathway (Fig. 5*A*) experiments where cholinergic receptor activation was induced with CCh application prior to WTET at EC-CA2. The field potential traces were normalized to the amplitude of the first fEPSP in the train of high-frequency tetanic stimuli. Fig. 6*B* depicts the integral of normalized tetanization (TET) area at EC-CA2 of single and two-pathway experiments carried out with and without CCh priming. A comparison of the total normalized TET area at EC-CA2 with and without CCh priming, as depicted in Fig. 6*C*, shows that there is a significant increase in the total normalized TET area at EC-CA2 after CCh priming in both single pathway (Fig. 6*C, U* test, *P* = 0.0205), and two-pathway experiments (Fig. 6*C, U* test, *P* = 0.0177) compared to their respective control experiments without CCh priming. This suggests that the priming effect by prior CCh-induced synaptic depression significantly increases the level of depolarization in CA2 neurons immediately in response to WTET at EC-CA2. This observed increase in TET area after CCh priming is also suggestive of a possible enhancement of transmitter release during tetanic stimulation in the CCh-primed experiments compared to control experiments without CCh priming, which may be partly contributing to the early phase of CCh-primed LTP at EC-CA2 observed in Fig. 3*A* and Fig. 5*A*. However, we noted from the paired pulse stimulation experiments in Fig. 5*C* that by 30 min into LTP induction, there is no significant presynaptic component in the reinforcement of LTP as observed by comparable levels of paired pulse ratios before and after tetanic stimulation at EC-CA2 in the two-pathway experiment.

**Fig. 6.**
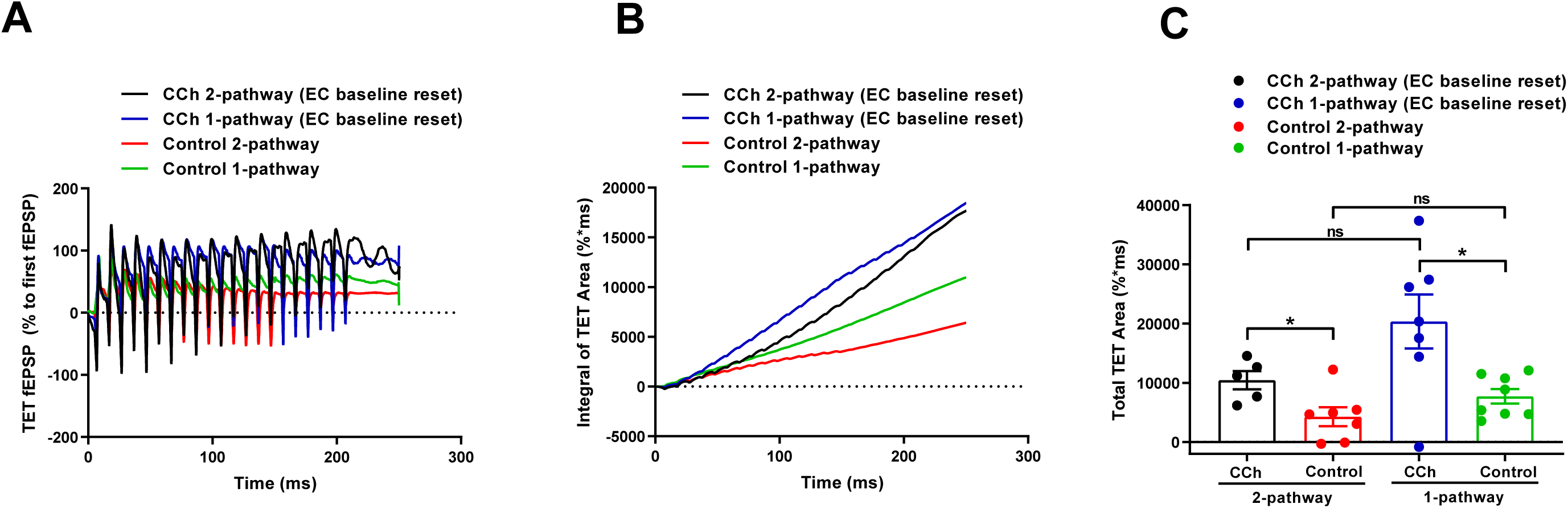
Comparison of the effect of CCh priming on tetanic stimuli mediated summation of field excitatory postsynaptic potentials at EC-CA2 synapses. *(A)* Average field potential traces in response to 100 Hz/s from single (*n* = 8; green line) and two-pathway (*n* = 7; red line) EC-CA2 WTET control experiments without CCh priming (from Fig. 3*B* and 5*B*, respectively), and from single (*n* = 7; blue line) and two-pathway (*n* = 5; black line) experiments with CCh-induced synaptic depression prior to WTET at EC-CA2 (from Fig. 3*A* and 5*A*, respectively). The field potential traces were normalized to the amplitude of the first fEPSP in the train of high frequency tetanic stimuli, and artefacts were truncated by linear interpolation. *(B)* Integral of normalized tetanization (TET) area at EC-CA2 of experiments carried out with and without CCh priming. *(C)* Comparison of the total normalized TET area of individual experiments, with and without CCh priming. Total normalized TET area at EC-CA2 after CCh priming in single and two-pathway experiments is significantly greater compared to their respective control experiments without CCh priming. Data show individual data points, mean ± SEM. Differences between groups are indicated as * *P* < 0.05 and ns *P* > 0.05.

Overall, our data demonstrate a cooperative interaction between concurrent CCh-induced LTD at SC-CA2 and the metaplastic reinforcement of LTP at EC-CA2 synapses. Interestingly, the CCh-primed LTP at EC-CA2 synapses is down-scaled by reversing the synaptic weights at SC-CA2 post CCh priming to baseline levels.

## Discussion

Cholinergic modulation of neuronal activity is critical to learning and memory processes in multiple neural systems (42, 43). Numerous studies deciphered the role of cholinergic signaling in the hippocampus for memory formation, particularly episodic and spatial memory, and synaptic plasticity, including LTP and LTD (44, 45). It has been shown that learning increases extracellular acetylcholine levels in the CA1 region of the hippocampus and injection of cholinergic antagonists into the CA1 region blocks learning-enhanced synaptic plasticity (46). However, few studies have explored the hippocampal subfield CA2 in this regard, barring the very recent study that has looked into cholinergic modulation of intrinsic properties and action potential firing of CA2 pyramidal neurons (41). Comprehensive studies pertaining to cholinergic modulation of long-term synaptic plasticity in the CA2 region were lacking. In this study, we demonstrate the dynamic modulation of synaptic plasticity in hippocampal area CA2 across time owing to cholinergic receptor activation with the non-selective cholinergic receptor agonist carbachol (CCh) that induces a protein synthesis-dependent long-term depression (CCh-LTD) at both the EC and SC synaptic inputs to CA2 neurons. While cholinergic receptor activation-mediated synaptic depression has been reported in multiple brain regions including hippocampal CA1 (47), CA3 (48), visual cortex (49), perirhinal cortex (50), and more recently in CA2 neurons (41), previous studies have not looked into the role of cholinergic modulation in the bidirectional and temporal regulation of plasticity at CA2 synapses that we have in the present study. The CCh-LTD at EC- and SC-CA2 synapses was also observed to be independent of both continuous afferent synaptic drive and NMDAR activation. This is in contrast to CCh application at the same concentration that induces a Hebbian form of LTD at CA3-CA1 synapses which is both activity and NMDAR dependent (47). However, like our results here, CCh-LTD in the perirhinal cortex (50) and medial prefrontal cortex (51) has been shown to independent of NMDAR activation and synaptic stimulation. This suggests that the mechanisms governing cholinergic receptor activation-induced synaptic depression varies between brain regions and hippocampal subfields. Our results further demonstrate that muscarinic M1, M2 and M3, and nicotinic ACh receptors are functionally present in area CA2, mediating different phases of CCh-LTD at EC- and SC-CA2 synapses. Of the three muscarinic receptors detected to play a role in CCh-LTD at CA2 synapses, activation of muscarinic M3 receptors appears to be most critical for the early induction of CCh-LTD.

Importantly, the present work demonstrates that prior induction of synaptic depression by cholinergic receptor activation acts as a ‘priming’ signal that subserves the metaplastic reinforcement of synaptic potentiation in CA2 neurons in response to a later bout of subthreshold plasticity-inducing stimuli. Metaplasticity regulates plasticity thresholds based on the history of synaptic activity such that modifications of neural activity at one point in time can alter the ability of synapses to subsequently generate synaptic plasticity such as LTP or LTD (52). We show that prior synaptic depression induced by cholinergic receptor activation primes EC-CA2 synapses for late LTP (L-LTP) in response to a later bout of weak tetanizing stimuli which normally elicits only a short-lasting early LTP (E-LTP). We therefore propose that metaplasticity triggered by priming activity of CCh-induced synaptic depression shifts the threshold (θ) for conventional strong tetanization induced L-LTP at EC-CA2 synapses to a lower threshold (θ^**’**^) such that weak tetanization can result in L-LTP, similar to a Bienenstock-Cooper-Munro model (BCM)-like sliding synaptic modification threshold that depends on the slowly varying time-averaged value of neural activity (53, 54). We also report a similar metaplastic effect at the SC inputs to CA2 neurons, such that CCh-induced prior synaptic depression allows for the induction and maintenance of L-LTP in response to strong tetanizing stimuli, which is otherwise ineffective at inducing LTP at the plasticity-resistant SC-CA2 synapses. Unlike the cortical inputs to CA2, the SC-CA2 inputs displayed only a transient potentiation in response to weak tetanic stimulation delivered subsequent to CCh-LTD at SC-CA2. This difference among EC- and SC-CA2 synapses in the strength of induction stimuli required for LTP reinforcement subsequent to CCh priming could be due to the inherent differences in plasticity expression along the proximo-distal dendritic axis of CA2 neurons, with the proximal SC-CA2 synapses being resistant to activity-dependent LTP. Nonetheless, strong tetanic stimuli delivered to SC-CA2 inputs following CCh ‘priming’ relieves this brake on synaptic potentiation and leads to the expression of L-LTP at SC-CA2.

Synaptic cross-tagging and cross-capture is an associative plasticity process wherein L-LTD/L-LTP at a synaptic input can help transform the protein synthesis-independent short-lasting form of the opposite plasticity process, namely E-LTP/E-LTD to their respective late forms at a neighbouring independent synaptic input (55) (56). This is because a late form of LTD or LTP at a synaptic input could possibly lead to *de novo* synthesis of a pool of plasticity-related proteins (PRPs) that may be common between LTD and LTP (process-independent PRPs) or a pool of process-dependent PRPs specific for LTD and LTP (55, 56). If LTP-specific PRPs or process-independent PRPs that could support both LTD and LTP, from this pool of PRPs synthesized downstream of L-LTD expressed at a synaptic input, are captured by the independent nearby synaptic input that is only weakly stimulated, it can lead to the transformation of E-LTP to L-LTP by virtue of the cross-capture (55, 56). Our results indicate that protein synthesis downstream of cholinergic stimulation and resultant CCh-LTD is required for the maintenance of LTP at the cortical inputs to CA2 neurons. Interestingly, protein synthesis inhibition at a later time point during the induction of E-LTP at EC-CA2 with weak tetanic stimulation does not abrogate the persistence of LTP, suggesting that it is not new protein synthesis in response to LTP-inducing stimuli, but the prior synthesis of proteins downstream of CCh-LTD that primes EC-CA2 synapses for L-LTP. This bears resemblance to the synaptic cross-tagging and capture process and suggests that a pool of process-independent PRPs common to LTD and LTP is likely synthesized at CA2 synapses during a period of high cholinergic tone allowing for the metaplastic reinforcement of E-LTP induced at a later time to L-LTP. It is possible that the activation of M1 and M3 muscarinic AChRs, and nicotinic AChRs during CCh-LTD plays a role in the observed metaplasticity in CA2 neurons conferred by cholinergic receptor activation. M1 and M3 mAChRs are G_q/11_ coupled receptors, downstream of which there is activation of the mitogen-activated protein kinase pathway extracellular signal-regulated kinases ERK1/2 (57) known to be involved in LTP and memory (58). ERK1/2 activation has also been shown to be critical for CCh-LTD (59, 60). It is possible that the upregulation of such PRPs downstream of cholinergic receptor activation could potentially pivot the synapses towards persistent LTP in response to subthreshold plasticity-inducing tetanizing stimuli. Muscarinic ACh receptor activation has also been shown to be involved in mediating the learning-induced synaptic delivery of AMPA receptors in CA1 neurons (46). Carbachol application has also been shown in a recent study to induce CA2 pyramidal neuron depolarization and burst firing of action potentials mediated by the activation of M1 and M3 mAChRs, and not M2 mAChRs (41). The G_q/11_-mediated signal downstream of M1, M3 mAChR activation can also mobilise intracellular calcium by Ca^2+^ release from inositol 1,4,5-trisphosphate (IP_3_)-sensitive intracellular stores, owing to phospholipase C activation (57), and the consequent rise in intracellular calcium levels can evoke signaling cascades that support LTP (61, 62). Likewise, nicotinic ACh receptors are ligand-gated cation permeable ion channels, some of which are permeable to calcium ions and it is possible that a resultant increase in cytoplasmic calcium levels may ensue signal transduction cascades (63) that could contribute to the observed CCh-primed LTP at CA2 synapses. Overall, it appears that a summation of multiple positive reinforcing effects in CA2 neurons downstream of activation of the cholinergic receptors implicated in CCh-LTD could serve to enhance the effectiveness of subthreshold plasticity-inducing stimuli to elicit lasting LTP, although the exact underlying mechanisms remain to be determined. We speculate a major role for M1 and M3 mAChR activation in lowering the threshold for LTP upon CCh priming to facilitate the metaplastic reinforcement of synaptic potentiation in CA2, given the prominent role for M1 and M3 receptors in the early phase of CCh-LTD manifest before reversing the stimulus strength toward baseline in the experiment timeline followed for the priming experiments in this study.

Our results also demonstrate a trans-compartmental regulation of synaptic weights along the proximo-distal dendritic axis of CA2 neurons such that CCh-primed LTP at distal cortical inputs to CA2 is modulated by varying strength of concurrent synaptic responses at the proximal Schaffer collateral pathway. We observed that while concurrent synaptic depression at SC-CA2 ‘cooperates’ with the stable expression of CCh-primed LTP at EC-CA2, reversing the synaptic weights at SC-CA2 toward baseline ‘interferes’ with the expression of primed LTP. This finding further underscores the importance of synaptic information processing between CA3 and CA2 in the expression of synaptic plasticity and memory mediated through the stimulation of cortical inputs to CA2 neurons. CA2, thus is not a mere connecting link between CA3 and CA1 but has an active role in the fate of information processing and storage in the cortico-hippocampal circuitry. That the CCh-primed LTP reinforcement is not displayed at EC-CA2 in the absence of sustained CCh-mediated synaptic depression at SC-CA2 in the two-pathway experiment favours the possibility that there exists a positive associative interaction between CCh-LTD at the Schaffer collateral inputs and the observed LTP at cortical inputs to CA2 neurons. What we observe in our study is that increasing synaptic activity at SC-CA2 following CCh-LTD does not lead to reinforcement of LTP at the heterosynaptic input EC-CA2, while the co-existence of CCh-LTD at SC-CA2 during delivery of a weak tetanic stimulus to EC-CA2 allows for the conversion of a transient E-LTP to L-LTP at EC-CA2. This may be regarded as a possible homeostatic mechanism to reset synaptic weights in accordance to synaptic activity patterns. Thus, metaplastic learning rules are extended to synaptic activity patterns between heterosynaptic pathways. Such metaplastic regulatory mechanisms can potentially facilitate normalization of synaptic weights (64, 65), and thereby ensure the stability of a dynamically learning neuronal network.

A depletion of cholinergic fibres in the CA2 region is observed in Parkinson’s disease (PD) cases with cognitive deficits and there exists a significant association between the high deposition of Lewy pathology in area CA2 and dementia in PD (66). It has also been observed in post-mortem brain samples of Alzheimer’s disease patients that the CA2 region displays a significant decrease in the density of ChAT-positive nerve fibres (29). The findings presented in this study should therefore prove relevant to our understanding of cognitive impairment in neurodegenerative disorders that feature cholinergic dysfunction such as Alzheimer’s disease (67) and Parkinson’s disease (68). The present work has focused largely on the regulation of synaptic plasticity at EC-CA2 synapses subsequent to the priming activity mediated by CCh-LTD. The mechanisms governing CCh-primed L-LTP observed at the plasticity-resistant SC-CA2 synapses merit future investigation. We conjecture that metaplastic synaptic modifications may significantly contribute to plasticity expression at the CA3→CA2 pathway when studied in the temporal context of neuronal activity, although conventional activity-dependent LTP is seemingly absent at these synapses. Together, the present study provides first evidence that priming activity by cholinergic receptor activation and the resultant synaptic depression in hippocampal CA2 neurons serves as a metaplastic trigger enabling a bidirectional switch to persistent LTP at EC- and SC-CA2 synapses. Hippocampal cholinergic tone is increased during active wakefulness observed in exploration of novel environments (69) and during spatial learning tasks (70). High hippocampal cholinergic tone during active wakefulness promotes theta oscillations that support memory encoding whereas low cholinergic tone during rest promotes sharp wave ripple oscillations that support memory consolidation (71-73). Thus, a dynamic modulation of CA2 cholinergic tone could potentially influence neuronal oscillations and thereby the encoding and subsequent consolidation of CA2-dependent memories. It is therefore plausible that a temporally sequenced modulation of synaptic strength at CA2 neurons subsequent to increased hippocampal cholinergic tone during exploratory behaviour, may contribute to the stability of encoded memories in a metaplastic manner. Indeed, sequential neuromodulation by first acetylcholine biases spike timing-dependent plasticity (STDP) toward depression at the Schaffer collateral-CA1 pathway, which when followed by the reward signal dopamine converts this synaptic depression to potentiation, and neural network models developed suggest that such sequential modulation of synaptic strength offers a mechanism for effective navigation and flexible learning in response to changing environments during reward-seeking foraging behaviour (74). Given that hippocampal CA2 is implicated in spatial, social and contextual memory (75-77), it would be of interest to study CA2 neuronal activity *in vivo* during exploratory behaviour and the effects of local manipulations of cholinergic receptors in the CA2 region and targeted activation or inactivation of basal forebrain cholinergic afferents to CA2, on CA2-dependent memory formation and behaviour. These outstanding possibilities will be of interest to explore in future studies.

## Materials and Methods

For electrophysiological experiments, acute hippocampal slices were prepared from young adult postnatal day (P)35-49 male Wistar rats. Animal procedures performed were approved by the guidelines of the Institutional Animal Care and Use Committee (IACUC) of the National University of Singapore. Detailed descriptions of hippocampal slice preparation, electrophysiology, pharmacology and statistical analysis are provided in ***SI Appendix***.

## Acknowledgments

We thank Dr Susan Jones (University of Cambridge, UK) for the constructive comments on the manuscript. This work was supported by National Medical Research Council Collaborative Research Grants (NMRC-CBRG-0099-2015 and NMRC/OFIRG/0037/2017), Ministry of Education Academic Research Fund Tier 3 (MOE2017-T3-1-002) and NUSMed-FoS Joint Research Programme (NUHSRO/2018/075/NUSMed-FoS/01) (to S.S.), and NUS Research Scholarship (to A.B.). T.B. was supported by the Natural Science Foundation of China (31871076) and Shanghai Municipal Science and Technology Major Project (No. 2018SHZDZX01) and ZJLab. The funder had no role in study design, data collection, or interpretation.

## Author contributions

A.B., T.B. and S.S. designed research. A.B. performed research. M.Z.B.I. assisted with some experiments. A.B., T.B. and S.S. analyzed data. A.B. and S.S. wrote the manuscript.

## Data Availability

All data that support the findings of this study are available upon request from the corresponding author.

## SI Figures

**Fig. S1.**
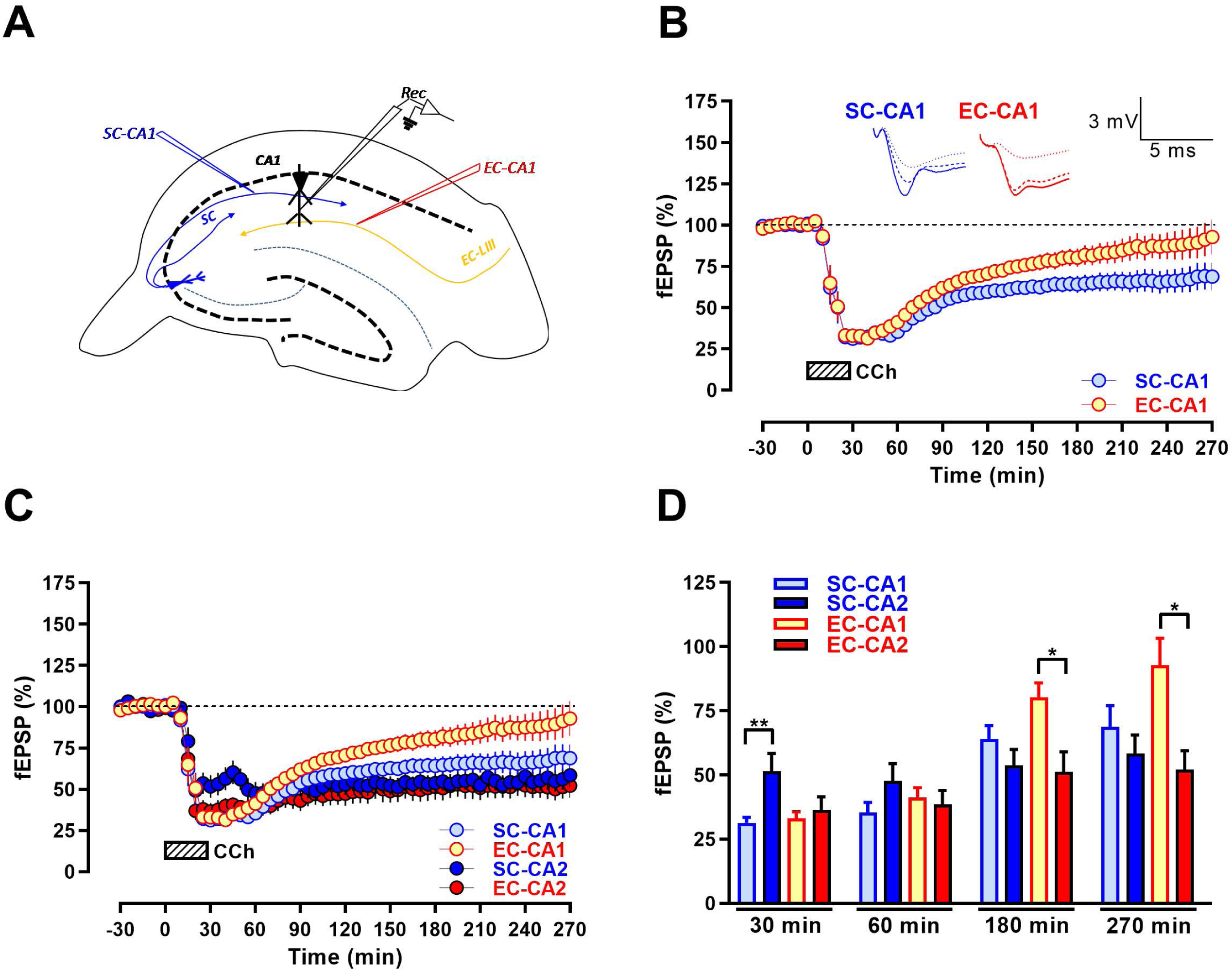
Comparison of CCh-induced depression between hippocampal CA1 and CA2 SR (SC) and SLM (EC) synapses. *(A)* A schematic diagram depicting the location of electrodes in the acute hippocampal slices for the extracellular field electrophysiology recordings from the CA1 region. Blue inverted triangle represents the placement of electrode in the *stratum radiatum* (SR) stimulating the Schaffer collateral (SC) afferents to CA1. Red inverted triangle represents the location of electrode in the *stratum lacunosum-moleculare* (SLM) stimulating the entorhinal cortical layer III (EC LIII) afferents to CA1. Black inverted triangle depicts the position of the recording electrode in the CA1 distal dendritic region. *(B)* Bath application of CCh (50 µM) for 30 min after a 30 min stable baseline induced a long-lasting synaptic depression at SC inputs to CA1, lasting up to 270 min into CCh application, while synaptic responses at EC inputs to CA1 reversed to its baseline levels by 210 min into CCh application (*n* = 10). *(C)* Superimposition of CCh-LTD at CA2 synapses (data from Fig. 1*C*) and CA1 synapses (data from Fig. S1*B*) for the ease of comparison. *(D)* Summary bar graph compares fEPSP(%) at 30 min, 60 min, 180 min and 270 min into CCh application at SC-CA1 and EC-CA1 synapses (from Fig. S1*B*) against SC-CA2 and EC-CA2 synapses (from Fig. 1*C*). All data show mean ± SEM. Significant differences between groups are indicated as * *P* < 0.05, ** *P* < 0.01. Analog traces in *B* depict representative SC-CA1 (blue) and EC-CA1 (red) fEPSPs within the initial 30 min baseline recording (closed line) before CCh application, and 60 min (dotted line) and 270 min (hatched line) after baseline. (Scale bar for analog traces in *B*: 3 mV/5 ms.)

**Fig. S2.**
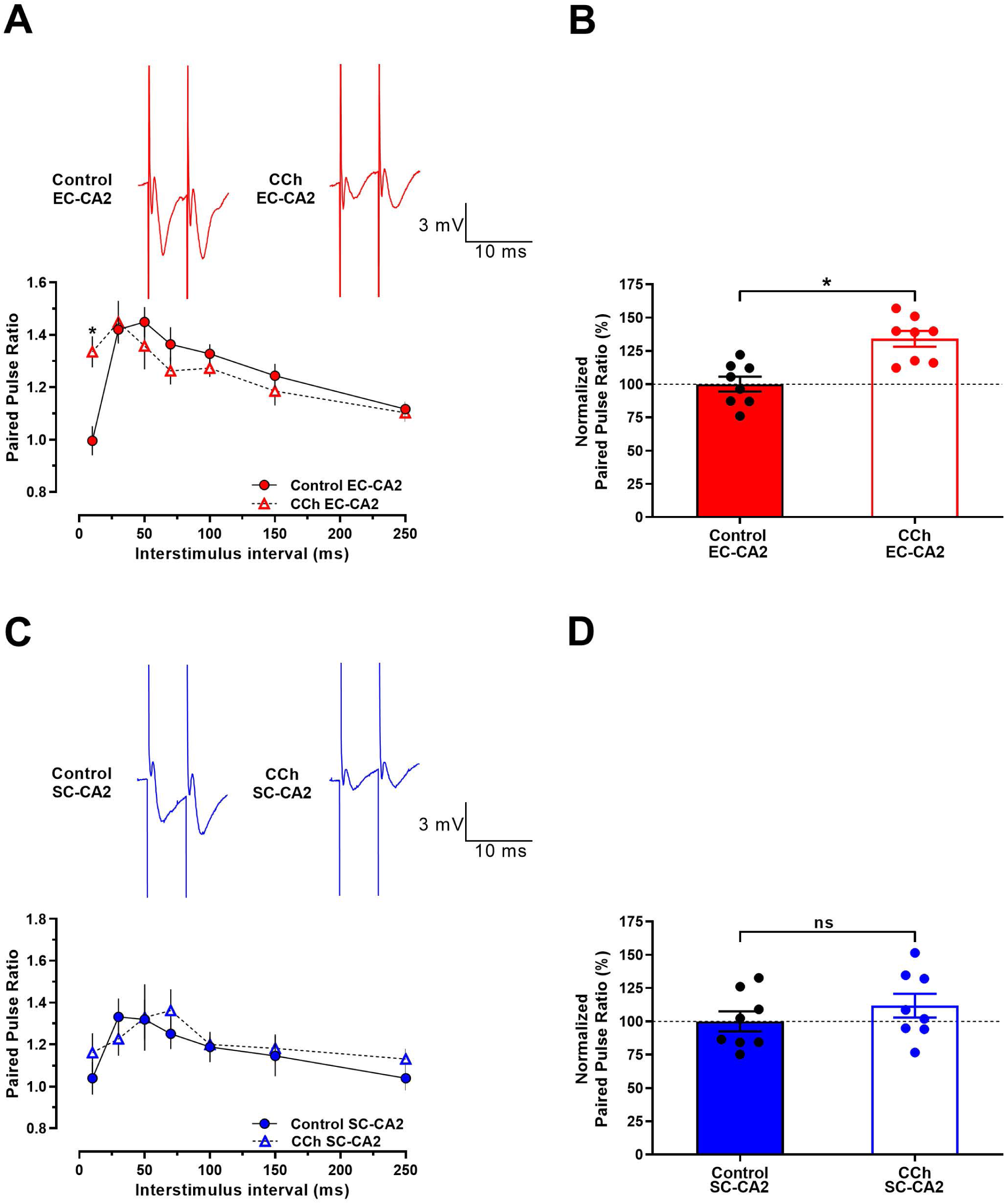
Cholinergic receptor activation differentially modulates presynaptic release probability at EC-CA2 and SC-CA2 synapses. *(A)* Paired pulse ratio at EC-CA2 before CCh application during the 30 min baseline recording (‘control’) and 60 min into 50 µM CCh application (‘CCh’) recorded for interstimulus intervals of 10 ms, 30 ms, 50 ms, 70 ms, 100 ms, 150 ms and 250 ms (*n* = 8). Paired pulse ratio at 10 ms interstimulus interval in EC-CA2 60 min into CCh application is significantly higher than control paired pulse ratio before CCh application. *(B)* Paired pulse ratio (%) at 10 ms interstimulus interval in EC-CA2 normalized by the average control paired pulse ratio shows a significant increase for 60 min into CCh application compared to control. *(C)* Paired pulse ratios at SC-CA2 were not significantly different before CCh (‘control’) and 60 min into CCh application (‘CCh’) at 10 ms, 30 ms, 50 ms, 70 ms, 100 ms, 150 ms and 250 ms interstimulus intervals (*n* = 8). *(D)* Paired pulse ratio (%) at 10 ms interstimulus interval in SC-CA2 normalized by the average control paired pulse ratio shows no significant change between control and CCh groups. All data show mean ± SEM. Differences between groups are indicated as * *P* < 0.05 and ns *P* > 0.05. Representative analog traces depict EC-CA2 (red) and SC-CA2 (blue) fEPSPs for the paired stimulation at 10 ms interstimulus interval. (Scale bars for analog traces in *A* and *C*: 3 mV/10 ms.)

**Fig. S3.**
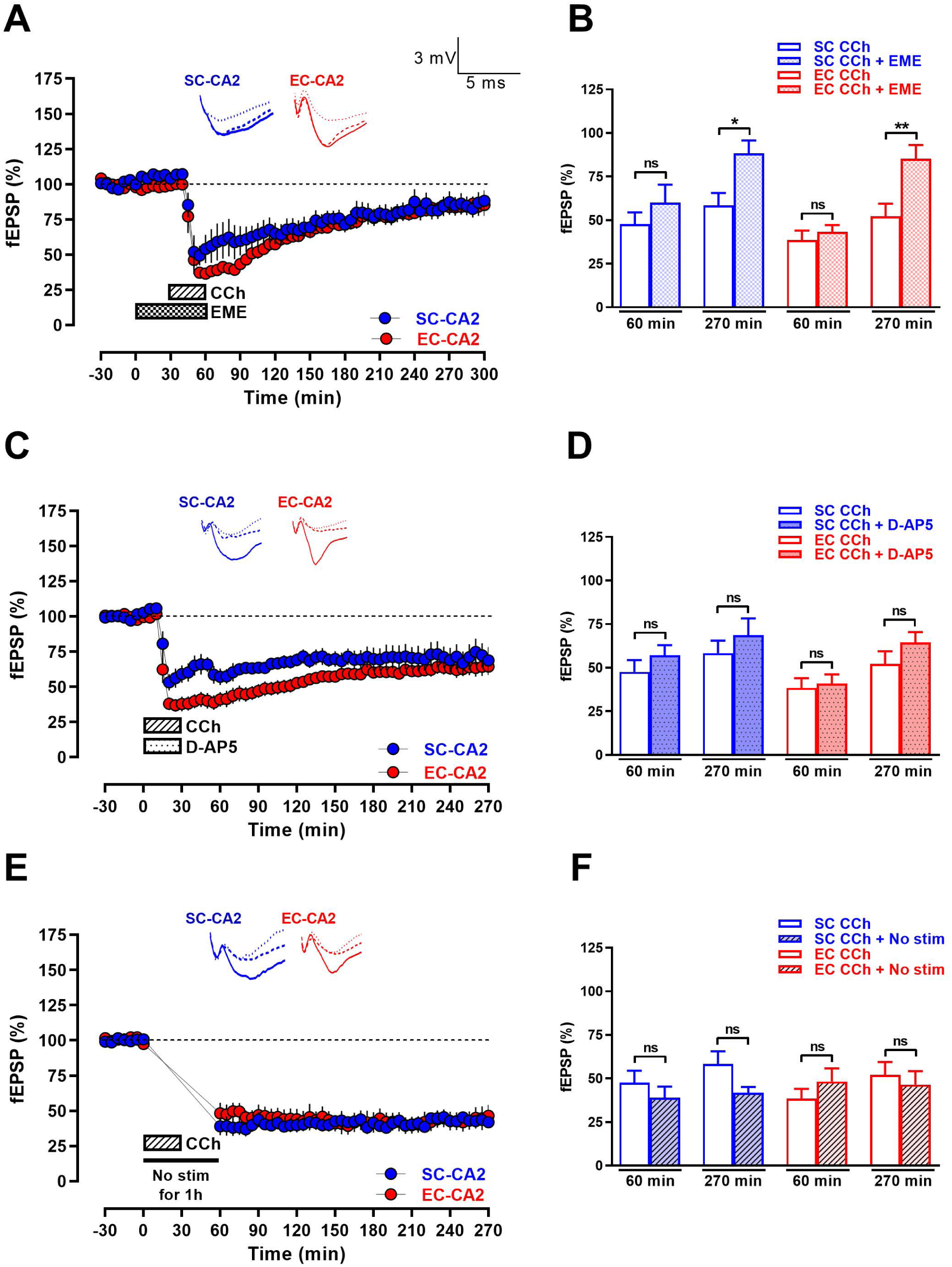
Persistence of CCh-LTD at CA2 synapses requires protein synthesis but is independent of NMDAR activation and continuous afferent synaptic drive. *(A)* The protein synthesis inhibitor, emetine (EME; 20 µM) is bath-applied for a total duration of 60 min (30 min before CCh and 30 min co-application with CCh) after a 30 min stable baseline, resulting in the blockade of the late phase of CCh-LTD at EC-CA2 and SC-CA2 (*n* = 7). *(B)* Summary bar graph comparing fEPSP(%) 60 min (acute CCh-LTD) and 270 min (sustained CCh-LTD) into CCh application between Fig. 1*C* (CCh alone) and Fig. S3*A* (CCh + emetine). *(C)* Co-application of the NMDAR antagonist D-AP5 (50 µM) with CCh for 30 min following a 30 min stable baseline does not abrogate CCh-LTD and manifests stable late CCh-LTD at EC-CA2 and SC-CA2 (*n* = 8). *(D)* Summary bar graph comparing fEPSP(%) 60 min and 270 min into CCh application between Fig. 1*C* (CCh alone) and Fig. S3*C* (CCh + D-AP5). *(E)* Suspension of basal test stimulus (‘No stim’) at EC-CA2 and SC-CA2 for 1 h (30 min during application of CCh and 30 min following CCh application) does not affect CCh-LTD and displays stable synaptic depression into the late phase (*n* = 6). *(F)* Summary bar graph comparing fEPSP(%) 60 min and 270 min into CCh application between Fig. 1*C* (CCh alone) and Fig. S3*E* (CCh + No stim). All data show mean ± SEM. Differences between groups are indicated as * *P* < 0.05, ** *P* < 0.01 and ns *P* > 0.05. Analog traces depict representative EC-CA2 (red) and SC-CA2 (blue) fEPSPs within 30 min before CCh application (closed line), at 60 min into CCh application (dotted line) and at 270 min into CCh application (hatched line). (Scale bars for analog traces in *A, C*, and *E*: 3 mV/5 ms.)

## Supplementary Information for

## Supplementary Information Text

### Materials and Methods

#### Animals and preparation of hippocampal slices

For electrophysiological experiments, a total of 215 acute hippocampal slices were used from 107 young adult male Wistar rats (P35-P49). All animals were kept under 12 h light/dark cycle with food and water available *ad libitum*. Rats were anesthetized using CO_2_ and thereafter decapitated, following which brains were removed and hippocampi quickly isolated in artificial cerebrospinal fluid (ACSF) maintained at 4°C and pH 7.3. The composition (in mM) of ACSF (1) is as follows: 124 NaCl, 2.5 KCl, 2 MgCl_2,_ 2 CaCl_2,_ 1.25 NaH_2_PO_4,_ 26 NaHCO_3_ and 17 D-Glucose, equilibrated with 95% O_2_ and 5% CO_2_ (carbogen; flow rate 16 L/h). Using a manual tissue chopper, transverse hippocampal slices with a thickness of 400 µm were prepared from the isolated hippocampi (right/left). Slices were then placed on a nylon net in the interphase chamber (Scientific System Design, Ontario, Canada) maintained at 32°C and preincubated for a minimum 2-3 h recovery period prior to placing electrodes. The ACSF flow rate to the chamber was maintained at 1 mL/min. Anesthetization, dissection and transfer of slices to the chamber were performed quickly and did not exceed an average duration of 5 min (2).

#### Extracellular field potential recording

Three monopolar lacquer-insulated stainless steel electrodes (5 MΩ; AM Systems, Sequim, WA, USA) were positioned in the hippocampal area CA2 for the two-pathway experiments, and two monopolar lacquer-insulated stainless steel electrodes were placed in area CA2 for the single-pathway experiments. Briefly, for the two-pathway experiments, two stimulating electrodes, one stimulating the Schaffer collateral (SC) fibres (CA3→CA2) and the other stimulating the entorhinal cortical fibres (EC LII→CA2) onto CA2, were positioned at the proximal and distal sites, respectively, and field excitatory postsynaptic potentials (fEPSP) upon stimulation of proximal and distal inputs were recorded from the CA2 distal dendritic region (Fig. 1*A*). The same applies for the single-pathway experiments performed in this study with the exception that in addition to the recording electrode at the CA2 distal dendritic region, only one stimulating electrode was positioned which stimulates either the proximal Schaffer collateral fibres or the distal entorhinal cortical fibres onto CA2. The recorded signals were amplified by a differential amplifier (Model 1700; AM Systems), digitized using a CED 1401 analog-to-digital converter (Cambridge Electronic Design, Cambridge, UK) and monitored on-line with Intracell software (IfN, Magdeburg, Germany) where the initial slope function (millivolts per millisecond) of the fEPSP is measured for analysis. The independence of the entorhinal cortical and Schaffer collateral synaptic inputs onto CA2 was confirmed using a cross-input paired pulse facilitation (PPF) protocol with an interstimulus interval of 50 ms similar to our previous study (1). The absence of PPF indicated that the two stimulating electrodes activated independent synaptic inputs (1). For the experiment in Fig. S1*B*, two stimulating electrodes, one stimulating the Schaffer collateral fibres (CA3→CA1) and the other stimulating the entorhinal cortical fibres (EC LIII→CA1) onto CA1, were positioned at the CA1 *stratum radiatum* and CA1 *stratum lacunosum-moleculare*, respectively, and fEPSPs were recorded from the CA1 distal dendritic region upon stimulation of the proximal SC and distal EC inputs (Fig. S1*A*).

After the preincubation period of 2-3 h for recovery of slices placed in the interphase chamber post dissection, an input-output relation (afferent stimulation intensity vs. fEPSP slope) for each stimulated synaptic input was generated to determine the test stimulation strength which was set to obtain an fEPSP response with 40% of the maximal fEPSP slope value. At this stimulus intensity, a stable baseline was recorded for 30 min before application of pharmacological agents or tetanizing stimuli. The basal stimulation consists of four sweeps of 0.2 Hz biphasic constant current pulses (pulse duration, 0.1 ms per polarity) at each time point (2). For early long-term potentiation (E-LTP) induction, a ‘weak’ tetanization (WTET) protocol consisting of a single train of high-frequency stimulation of 21 pulses at 100 Hz (pulse duration, 0.2 ms per polarity) was delivered (2). The ‘strong’ tetanization (STET) protocol consisted of three trains of high-frequency stimulation of 100 pulses at 100 Hz (pulse duration, 0.2 ms per polarity) with an inter-train interval of 10 min (2). Paired pulse stimulation experiments in Fig. S2*A*, Fig. S2*C* and Fig. 5*C* were performed by delivering pairs of stimulation pulses with an interstimulus interval of 10 ms, 30 ms, 50 ms, 70 ms, 100 ms, 150 ms and 250 ms, to individual synaptic inputs at specified time points (described in the results section and figure legends) of the fEPSP recording for the respective experiments. All field potential recordings in this study were performed in the absence of application of GABA receptor antagonists similar to our previous studies (1, 3), and therefore reflect responses from hippocampal neurons with intact inhibitory transmission.

#### Pharmacology

The non-selective cholinergic receptor agonist, (2-Hydroxyethyl)trimethylammonium chloride carbamate (carbachol; C4382; Sigma-Aldrich), was prepared as 50 mM stock in deionized water and diluted to a final concentration of 50 µM in ACSF. Protein synthesis inhibitor, emetine dihydrochloride hydrate (emetine; E2375; Sigma-Aldrich) was prepared as 20 mM stock in deionized water and diluted to 20 µM final concentration in ACSF. The N-methyl-D-aspartate receptor (NMDAR) antagonist, D-2-amino-5-phosphonovalerate (D-AP5; 0106; Tocris Bioscience), was stored at -20°C as 50 mM stock prepared in deionized water and diluted before application to a final concentration of 50 µM in ACSF. The M1 muscarinic receptor antagonist, 5,11-Dihydro-11-[(4-methyl-1-piperazinyl)acetyl]-6*H*-pyrido[2,3-*b*][1,4] benzodiazepin-6-one dihydrochloride (pirenzepine; 1071; Tocris Bioscience) was prepared as 10 mM stock in deionized water and diluted to a final concentration of 0.5 µM in ACSF. The M2 muscarinic receptor antagonist, 11-[[2-[(Diethylamino)methyl]-1-piperidinyl]acetyl]-5,11-dihydro-6*H*-pyrido[2,3-*b*][1,4]benzodiazepin-6-one (AF-DX 116; 1105; Tocris Bioscience) was prepared as 10 mM stock in dimethyl sulfoxide (DMSO), stored at -20°C and diluted before application to a final concentration of 2 µM in ACSF. The M3 muscarinic receptor antagonist, 1,1-Dimethyl-4-diphenylacetoxypiperidinium iodide (4-DAMP; 0482; Tocris Bioscience), was prepared as 10 mM stock in DMSO, stored at -20°C and diluted before application to a final concentration of 1 µM in ACSF. The non-selective muscarinic receptor antagonist, (endo,syn)-(±)-3-(3-Hydroxy-1-oxo-2-phenylpropoxy)-8-methyl-8-(1-methylethyl)-8-azoniabicyclo[3.2.1]octane bromide (atropine; 0692; Tocris Bioscience) was prepared as 5 mM stock in deionized water and diluted to a final concentration of 1 µM in ACSF. The non-competitive nicotinic receptor antagonist, *N*,2,3,3-Tetramethylbicyclo[2.2.1]heptan-2-amine hydrochloride (mecamylamine; 2843; Tocris Bioscience) was prepared as 20 mM stock in deionized water and diluted to a final concentration of 20 µM in ACSF. The concentrated pharmacological agent stock solutions were diluted in ACSF to their respective final concentrations shortly before application, after which the pharmacological agent-containing ACSF was saturated with carbogen and bath-applied. Detailed descriptions of the duration and time point of application for individual pharmacological agents are provided in the results section. Where DMSO was used to prepare the stock solution, the final DMSO concentration in the working solution was kept at or below 0.1%, a concentration shown in previous studies to not affect basal synaptic transmission (4). Light-sensitive pharmacological agents were shielded from exposure to light during storage and bath application.

#### Statistical Analysis

fEPSP values are represented as the mean of normalized fEPSP slope function (percentage of baseline) ± SEM. When compared within the same group, mean normalized fEPSPs were analysed using the Wilcoxon signed-rank test (Wilcox test). When compared between groups, the values were analysed using the Mann-Whitney *U* test (*U* test). Kruskal-Wallis test with Dunn’s multiple comparison *post hoc* analysis was used to compare paired pulse ratios between three groups. Kolmogorov-Smirnov test was used to compare cumulative frequency distributions. Differences at *P* < 0.05 were considered statistically significant (* *P* < 0.05, ** *P* < 0.01, *\*\*\* P* < 0.001, *\*\*\*\* P* < 0.0001). Nonparametric tests were used as the sample size per series did not always guarantee a Gaussian normal distribution of the data. Prism version 8.0 (GraphPad Software, San Diego, CA) was used to plot the graphs and for the statistical analysis.

## References

1. J. Palacios-Filardo, J. R. Mellor, Neuromodulation of hippocampal long-term synaptic plasticity. Curr Opin Neurobiol 54, 37–43 (2019).

2. E. Marder, Neuromodulation of neuronal circuits: back to the future. Neuron 76, 1–11 (2012).

3. M. Aly, N. B. Turk-Browne, “How Hippocampal Memory Shapes, and Is Shaped by, Attention” in The Hippocampus from Cells to Systems: Structure, Connectivity, and Functional Contributions to Memory and Flexible Cognition, D. E. Hannula, M. C. Duff, Eds. (Springer International Publishing, Cham, 2017), 10.1007/978-3-319-50406-3_12, pp. 69–403.

4. C. G. Kentros, N. T. Agnihotri, S. Streater, R. D. Hawkins, E. R. Kandel, Increased attention to spatial context increases both place field stability and spatial memory. Neuron 42, 283–295 (2004).

5. J. Josh Lawrence, S. Cobb, “Neuromodulation of Hippocampal Cells and Circuits” in Hippocampal Microcircuits: A Computational Modeler’s Resource Book, V. Cutsuridis, B. P. Graham, S. Cobb, I. Vida, Eds. (Springer International Publishing, Cham, 2018), 10.1007/978-3-319-99103-0_7, pp. 227–325.

6. A. Benoy, A. Dasgupta, S. Sajikumar, Hippocampal area CA2: an emerging modulatory gateway in the hippocampal circuit. Exp Brain Res 236, 919–931 (2018).

7. H. J. Lee, H. K. Caldwell, A. H. Macbeth, S. G. Tolu, W. S. Young, 3rd, A conditional knockout mouse line of the oxytocin receptor. Endocrinology 149, 3256–3263 (2008).

8. W. S. Young, J. Li, S. R. Wersinger, M. Palkovits, The vasopressin 1b receptor is prominent in the hippocampal area CA2 where it is unaffected by restraint stress or adrenalectomy. Neuroscience 143, 1031–1039 (2006).

9. S. M. Dudek, G. M. Alexander, S. Farris, Rediscovering area CA2: unique properties and functions. Nat Rev Neurosci 17, 89–102 (2016).

10. Z. Borhegyi, C. Leranth, Substance P innervation of the rat hippocampal formation. J Comp Neurol 384, 41–58 (1997).

11. S. Chen et al., A hypothalamic novelty signal modulates hippocampal memory. Nature 586, 270–274 (2020).

12. V. Robert et al., Local circuit allowing hypothalamic control of hippocampal area CA2 activity and consequences for CA1. bioRxiv (2020).

13. Z. Cui, C. R. Gerfen, W. S. Young, 3rd, Hypothalamic and other connections with dorsal CA2 area of the mouse hippocampus. J Comp Neurol 521, 1844–1866 (2013).

14. K. Yoshida, H. Oka, Topographical projections from the medial septum-diagonal band complex to the hippocampus: a retrograde tracing study with multiple fluorescent dyes in rats. Neurosci Res 21, 199–209 (1995).

15. R. C. Meibach, A. Siegel, Efferent connections of the septal area in the rat: An analysis utilizing retrograde and anterograde transport methods. Brain Research 119, 1–20 (1977).

16. C. Nyakas, P. G. Luiten, D. G. Spencer, J. Traber, Detailed projection patterns of septal and diagonal band efferents to the hippocampus in the rat with emphasis on innervation of CA1 and dentate gyrus. Brain Res Bull 18, 533–545 (1987).

17. R. P. Gaykema, J. van der Kuil, L. B. Hersh, P. G. Luiten, Patterns of direct projections from the hippocampus to the medial septum-diagonal band complex: anterograde tracing with Phaseolus vulgaris leucoagglutinin combined with immunohistochemistry of choline acetyltransferase. Neuroscience 43, 349–360 (1991).

18. V. Chevaleyre, S. A. Siegelbaum, Strong CA2 pyramidal neuron synapses define a powerful disynaptic cortico-hippocampal loop. Neuron 66, 560–572 (2010).

19. K. Kohara et al., Cell type-specific genetic and optogenetic tools reveal hippocampal CA2 circuits. Nat Neurosci 17, 269–279 (2014).

20. R. Bartesaghi, T. Gessi, Parallel activation of field CA2 and dentate gyrus by synaptically elicited perforant path volleys. Hippocampus 14, 948–963 (2004).

21. M. Zhao, Y. S. Choi, K. Obrietan, S. M. Dudek, Synaptic plasticity (and the lack thereof) in hippocampal CA2 neurons. J Neurosci 27, 12025–12032 (2007).

22. S. E. Lee et al., RGS14 is a natural suppressor of both synaptic plasticity in CA2 neurons and hippocampal-based learning and memory. Proc Natl Acad Sci U S A 107, 16994–16998 (2010).

23. P. R. Evans et al., RGS14 Restricts Plasticity in Hippocampal CA2 by Limiting Postsynaptic Calcium Signaling. eNeuro 5 (2018).

24. S. B. Simons, Y. Escobedo, R. Yasuda, S. M. Dudek, Regional differences in hippocampal calcium handling provide a cellular mechanism for limiting plasticity. Proc Natl Acad Sci U S A 106, 14080–14084 (2009).

25. K. E. Carstens, M. L. Phillips, L. Pozzo-Miller, R. J. Weinberg, S. M. Dudek, Perineuronal Nets Suppress Plasticity of Excitatory Synapses on CA2 Pyramidal Neurons. J Neurosci 36, 6312–6320 (2016).

26. A. Dasgupta et al., Group III metabotropic glutamate receptors gate long-term potentiation and synaptic tagging/capture in rat hippocampal area CA2. Elife 9 (2020).

27. A. Dasgupta et al., Substance P induces plasticity and synaptic tagging/capture in rat hippocampal area CA2. Proc Natl Acad Sci U S A 114, E8741–E8749 (2017).

28. J. H. Pagani et al., Role of the vasopressin 1b receptor in rodent aggressive behavior and synaptic plasticity in hippocampal area CA2. Mol Psychiatry 20, 490–499 (2015).

29. G. Ransmayr et al., Choline acetyltransferase-like immunoreactivity in the hippocampal formation of control subjects and patients with Alzheimer’s disease. Neuroscience 32, 701–714 (1989).

30. E. Orta-Salazar, C. A. Cuellar-Lemus, S. Diaz-Cintra, A. I. Feria-Velasco, Cholinergic markers in the cortex and hippocampus of some animal species and their correlation to Alzheimer’s disease. Neurologia 29, 497–503 (2014).

31. T. Ichikawa, Y. Hirata, Organization of choline acetyltransferase-containing structures in the forebrain of the rat. J Neurosci 6, 281–292 (1986).

32. S. I. Mellgren, W. Harkmark, B. Srebro, Some enzyme histochemical characteristics of the human hippocampus. Cell Tissue Res 181, 459–471 (1977).

33. I. Bakst, D. G. Amaral, The distribution of acetylcholinesterase in the hippocampal formation of the monkey. J Comp Neurol 225, 344–371 (1984).

34. W. C. Abraham, M. F. Bear, Metaplasticity: the plasticity of synaptic plasticity. Trends Neurosci 19, 126–130 (1996).

35. W. C. Abraham, W. P. Tate, Metaplasticity: a new vista across the field of synaptic plasticity. Prog Neurobiol 52, 303–323 (1997).

36. M. E. Hasselmo, E. Schnell, Laminar selectivity of the cholinergic suppression of synaptic transmission in rat hippocampal region CA1: computational modeling and brain slice physiology. J Neurosci 14, 3898–3914 (1994).

37. D. A. Brown, Acetylcholine and cholinergic receptors. Brain Neurosci Adv 3, 2398212818820506 (2019).

38. A. I. Levey, S. M. Edmunds, V. Koliatsos, R. G. Wiley, C. J. Heilman, Expression of m1-m4 muscarinic acetylcholine receptor proteins in rat hippocampus and regulation by cholinergic innervation. J Neurosci 15, 4077–4092 (1995).

39. P. Seguela, J. Wadiche, K. Dineley-Miller, J. A. Dani, J. W. Patrick, Molecular cloning, functional properties, and distribution of rat brain alpha 7: a nicotinic cation channel highly permeable to calcium. J Neurosci 13, 596–604 (1993).

40. E. Wada et al., Distribution of alpha 2, alpha 3, alpha 4, and beta 2 neuronal nicotinic receptor subunit mRNAs in the central nervous system: a hybridization histochemical study in the rat. J Comp Neurol 284, 314–335 (1989).

41. V. Robert et al., The mechanisms shaping CA2 pyramidal neuron action potential bursting induced by muscarinic acetylcholine receptor activation. J Gen Physiol 152 (2020).

42. M. E. Hasselmo, The role of acetylcholine in learning and memory. Curr Opin Neurobiol 16, 710–715 (2006).

43. P. E. Gold, Acetylcholine modulation of neural systems involved in learning and memory. Neurobiol Learn Mem 80, 194–210 (2003).

44. J. Haam, J. L. Yakel, Cholinergic modulation of the hippocampal region and memory function. J Neurochem 142 Suppl 2, 111–121 (2017).

45. L. M. Teles-Grilo Ruivo, J. R. Mellor, Cholinergic modulation of hippocampal network function. Front Synaptic Neurosci 5, 2 (2013).

46. D. Mitsushima, A. Sano, T. Takahashi, A cholinergic trigger drives learning-induced plasticity at hippocampal synapses. Nat Commun 4, 2760 (2013).

47. C. L. Scheiderer et al., Sympathetic sprouting drives hippocampal cholinergic reinnervation that prevents loss of a muscarinic receptor-dependent long-term depression at CA3-CA1 synapses. J Neurosci 26, 3745–3756 (2006).

48. S. Williams, D. Johnston, Muscarinic depression of synaptic transmission at the hippocampal mossy fiber synapse. J Neurophysiol 64, 1089–1097 (1990).

49. A. Kirkwood, C. Rozas, J. Kirkwood, F. Perez, M. F. Bear, Modulation of long-term synaptic depression in visual cortex by acetylcholine and norepinephrine. J Neurosci 19, 1599–1609 (1999).

50. P. V. Massey, G. Bhabra, K. Cho, M. W. Brown, Z. I. Bashir, Activation of muscarinic receptors induces protein synthesis-dependent long-lasting depression in the perirhinal cortex. Eur J Neurosci 14, 145–152 (2001).

51. D. A. Caruana, E. C. Warburton, Z. I. Bashir, Induction of activity-dependent LTD requires muscarinic receptor activation in medial prefrontal cortex. J Neurosci 31, 18464–18478 (2011).

52. W. C. Abraham, Metaplasticity: tuning synapses and networks for plasticity. Nat Rev Neurosci 9, 387 (2008).

53. E. L. Bienenstock, L. N. Cooper, P. W. Munro, Theory for the development of neuron selectivity: orientation specificity and binocular interaction in visual cortex. J Neurosci 2, 32–48 (1982).

54. P. Jedlicka, Synaptic plasticity, metaplasticity and BCM theory. Bratisl Lek Listy 103, 137–143 (2002).

55. S. Frey, J. U. Frey, ‘Synaptic tagging’ and ‘cross-tagging’ and related associative reinforcement processes of functional plasticity as the cellular basis for memory formation. Prog Brain Res 169, 117–143 (2008).

56. S. Sajikumar, J. U. Frey, Late-associativity, synaptic tagging, and the role of dopamine during LTP and LTD. Neurobiol Learn Mem 82, 12–25 (2004).

57. A. A. Lanzafame, A. Christopoulos, F. Mitchelson, Cellular signaling mechanisms for muscarinic acetylcholine receptors. Recept Channels 9, 241–260 (2003).

58. S. Impey, K. Obrietan, D. R. Storm, Making new connections: role of ERK/MAP kinase signaling in neuronal plasticity. Neuron 23, 11–14 (1999).

59. C. L. Scheiderer et al., Coactivation of M(1) muscarinic and alpha1 adrenergic receptors stimulates extracellular signal-regulated protein kinase and induces long-term depression at CA3-CA1 synapses in rat hippocampus. J Neurosci 28, 5350–5358 (2008).

60. P. A. McCoy, L. L. McMahon, Muscarinic receptor dependent long-term depression in rat visual cortex is PKC independent but requires ERK1/2 activation and protein synthesis. J Neurophysiol 98, 1862–1870 (2007).

61. D. Fernandez de Sevilla, A. Nunez, M. Borde, R. Malinow, W. Buno, Cholinergic-mediated IP3-receptor activation induces long-lasting synaptic enhancement in CA1 pyramidal neurons. J Neurosci 28, 1469–1478 (2008).

62. N. E. Karagas, K. Venkatachalam, Roles for the Endoplasmic Reticulum in Regulation of Neuronal Calcium Homeostasis. Cells 8 (2019).

63. J. L. Yakel, Cholinergic receptors: functional role of nicotinic ACh receptors in brain circuits and disease. Pflugers Arch 465, 441–450 (2013).

64. K. D. Miller, Synaptic economics: competition and cooperation in synaptic plasticity. Neuron 17, 371–374 (1996).

65. P. Yger, M. Gilson, Models of Metaplasticity: A Review of Concepts. Front Comput Neurosci 9, 138 (2015).

66. A. K. L. Liu et al., Hippocampal CA2 Lewy pathology is associated with cholinergic degeneration in Parkinson’s disease with cognitive decline. Acta Neuropathol Commun 7, 61 (2019).

67. E. J. Mufson, S. E. Counts, S. E. Perez, S. D. Ginsberg, Cholinergic system during the progression of Alzheimer’s disease: therapeutic implications. Expert Rev Neurother 8, 1703–1718 (2008).

68. S. Perez-Lloret, F. J. Barrantes, Deficits in cholinergic neurotransmission and their clinical correlates in Parkinson’s disease. NPJ Parkinsons Dis 2, 16001 (2016).

69. M. G. Giovannini et al., Effects of novelty and habituation on acetylcholine, GABA, and glutamate release from the frontal cortex and hippocampus of freely moving rats. Neuroscience 106, 43–53 (2001).

70. F. Fadda, S. Cocco, R. Stancampiano, Hippocampal acetylcholine release correlates with spatial learning performance in freely moving rats. Neuroreport 11, 2265–2269 (2000).

71. M. E. Hasselmo, J. McGaughy, High acetylcholine levels set circuit dynamics for attention and encoding and low acetylcholine levels set dynamics for consolidation. Prog Brain Res 145, 207–231 (2004).

72. M. Vandecasteele et al., Optogenetic activation of septal cholinergic neurons suppresses sharp wave ripples and enhances theta oscillations in the hippocampus. Proc Natl Acad Sci U S A 111, 13535–13540 (2014).

73. G. Buzsaki, Two-stage model of memory trace formation: a role for “noisy” brain states.Neuroscience 31, 551–570 (1989).

74. Z. Brzosko, S. Zannone, W. Schultz, C. Clopath, O. Paulsen, Sequential neuromodulation of Hebbian plasticity offers mechanism for effective reward-based navigation. Elife 6 (2017).

75. G. M. Alexander et al., Social and novel contexts modify hippocampal CA2 representations of space. Nat Commun 7, 10300 (2016).

76. F. L. Hitti, S. A. Siegelbaum, The hippocampal CA2 region is essential for social memory. Nature 508, 88–92 (2014).

77. M. E. Wintzer, R. Boehringer, D. Polygalov, T. J. McHugh, The hippocampal CA2 ensemble is sensitive to contextual change. J Neurosci 34, 3056–3066 (2014).

## SI References

1. A. Dasgupta et al., Substance P induces plasticity and synaptic tagging/capture in rat hippocampal area CA2. Proc Natl Acad Sci U S A 114, E8741–E8749 (2017).

2. M. S. Shetty et al., Investigation of Synaptic Tagging/Capture and Cross-capture using Acute Hippocampal Slices from Rodents. J Vis Exp 10.3791/53008 (2015).

3. A. Dasgupta et al., Group III metabotropic glutamate receptors gate long-term potentiation and synaptic tagging/capture in rat hippocampal area CA2. Elife 9 (2020).

4. S. Navakkode, S. Sajikumar, J. U. Frey, The type IV-specific phosphodiesterase inhibitor rolipram and its effect on hippocampal long-term potentiation and synaptic tagging. J Neurosci 24, 7740–7744 (2004).

